# Time in shells: Complex interaction between biological clock and biomineralisation in *Mytilus galloprovincialis*

**DOI:** 10.64898/2026.02.23.707208

**Authors:** Victoria Louis, Erwan Peru, Charles-Hubert Paulin, Franck Lartaud, Laurence Besseau

**Author notes:** co-last authors (Franck Lartaud, Laurence Besseau).

## Abstract

The growth of bivalve shells is neither homogeneous nor continuous in time, resulting in the formation of growth patterns that correspond to the alternation of growth lines and increments deposited at regular intervals of time. The control of periodic increment formation is poorly understood and several hypotheses have been proposed. It has been proposed that environmental factors directly impact shell growth patterns, although it occasionally fails to adequately explain the observed shell growth patterns. The present study investigates the alternative hypothesis that the process of shell biomineralisation is controlled by biological clocks. This study demonstrates the existence of a functional circadian clock in *M. galloprovincialis*, as evidenced by molecular and behavioural results. Core circadian clock genes and biomineralisation genes have been observed to be expressed in the same cells of the mantle as revealed by *in situ* hybridisation experiments. However, the expression of core circadian clock genes and biomineralisation genes tested *in situ* and *in aquaria* exhibited different rhythmic profiles. This finding suggests that the clock does not directly activate the expression of the targeted biomineralisation genes in the mantle. Nevertheless, a significant rhythm of expression of biomineralisation-related genes was observed in mussels reared under free-running conditions, revealing the endogenous nature of the rhythm. The present study suggests that biological clocks play a role in controlling shell biomineralisation in *M. galloprovincialis*, although the precise underlying mechanism remains to be elucidated.

## Introduction

Bivalve shells are complex structures composed of calcium carbonate and organic compounds[1]. Although composing only 0.1 to 5% of the total weight of the shell, organic compounds have a role in stabilising the unstable nucleus of CaCO_3_ and is involved in controlling crystal growth and organisation [2,3]. The biomineralisation process by which the shell is secreted is neither homogeneous nor continuous in time, leading to the formation of growth patterns [4]. In bivalve shells they consist of the alternation of growth lines and increments. Depending on the environmental context, increments are mainly formed on a daily (*i.e*., ∼24 hours) or tidal (*i.e*., ∼12.4 hours) basis. Other periodicities such as annual (*i.e*., ∼365.25 days), circalunar (*i.e*., ∼29.5 days) and circalunidian (*i.e*., ∼24.8 hours) are embedded in bivalve shells [5]. Those cycles are related to environmental variations linked to the interactions between Earth-Moon and Sun. However, the number of increments observed does not *sensu-stricto* reflect environmental periodic variations. In *Mytilus galloprovincialis*, a mussel species endemic to the Mediterranean region characterised by its absence of tides, the anticipated daily (*i.e*., ∼1 increment per day) or tidal increments (*i.e*., ∼ 2 increments per day) are not observed, but rather ∼1.7 increments per day [6], thus questioning what generates such a rhythm of shell biomineralisation. Interestingly while bivalves are reared in constant environmental conditions, they still form increments at regular intervals of time [7,8]. Those observations led to the hypothesis that biological clocks might be involved in the control of periodic increment formation in the shell of bivalves [4–6,8].

Biological clocks are auto-regulatory transcriptional and translational feedback loops (TTFLs) that are known to synchronise biological activities and allow organisms to anticipate environmental variability [9,10]. A hallmark of biological clocks is that they are endogenous processes, meaning the clock keeps ticking under constant environmental conditions (*i.e*., under free-running conditions) [11]. In bivalves, a theorical clock mechanism has been proposed in the oyster *Magallana gigas* based on the rhythmic gene expression of circadian clock molecular components [12]. In this TTFL, the CLOCK:BMAL1 complex bind to an E-Box, activating the transcription of its own repressor, *period, cryptochrome 2* and *timeless* genes. A coupled feedback loop consisting of the genes *Rorb* and *Reverb* regulates the expression of *Bmal1* [12,13]. The molecular overlap between the circadian and the circatidal clock is still debated. Recently, it has been shown that the positive actor, BMAL1 is shared between the two clocks in crustaceans [14,15]. In *M. gigas*, the most core components of the circadian clock have been shown to run tidal after entrainment to tides [16].

Interestingly in *M. galloprovincialis*, it has been shown that the rhythm of expression of genes implicated in the biomineralisation process reflects the incrementation pattern within the shell [6]. Consequently, it is tempting to hypothesise that a biological clock could regulate the transcription of genes involved in shell biomineralisation in this species. Subsequently, in this study, we tested the hypothesis that the rhythmicity of biomineralisation observed in the shell is controlled by the biological clock. To do this, we used the Mediterranean mussel *M. galloprovincialis* as a biological model. Our recent work on *M. galloprovincialis* sampled in Mediterranean lagunas, *i.e*. an environment where tidal signals are weak, suggests that the growth pattern observed in their shell may reflect an interaction between the environment and the biological clock [6]. In the present study, the putative components of a biological clock were identified in the *M. galloprovincialis* genome and their gene expression rhythms were assessed in specimens sampled at sea. The endogenous nature of circadian and circatidal rhythms in *M. galloprovincialis* were assessed in constant conditions using the valve activity [17]. Then, the rhythmic expression of both clock and biomineralisation genes was characterised in controlled environmental conditions. Two environmental cues were used to test the rhythmic gene expression: the light as it is the main time giver for the circadian clock [18] and the food availability as it has been shown to be a crucial factor for biomineralisation and a known time giver of the circadian clock [19–21].

## Material and Methods

### 1. Model species, study site and rearing protocols

The mussels (*Mytilus galloprovincialis*) used in this study originated from Mediterranean Sea on the sea shore of Banyuls-sur-Mer (Occitany, France). They were sampled between November 2019 and November 2021 at a depth of between 1 and 3 m on the anchor chain of the SOLA buoy, an autonomous station monitoring the oceanographic parameters of the bay of Banyuls-sur-Mer, belonging to the Oceanological Observatory of Banyuls-sur-Mer (OOB) and part of the ILICO and JERICO national and European research infrastructures.

#### a. Study site at sea

Ten mussels were collected every four hours at the SOLA site (GPS: 42° 29’ 300N, 03° 08’ 700E) over a period of 36 hours, from noon on 9 September 2020 to noon on 11 September 2020. A piece of the posterior region of the mantle was sampled and flash-frozen in liquid nitrogen for molecular analysis. Night samples were taken under red light. Tides timetables were obtained on the Service national d’Hydrographie et d’Océanographie de la Marine (SHOM) website [22].

#### b. Mussel rearing under controlled environmental conditions

Mussels were reared in 36 L aquaria supplied with natural seawater by a semi-open system installed in a thermoregulated room at 16°C. The water was pumped from the bay of Banyuls-sur-Mer and pre-filtered at 200 µm before being stored in a buffer tank. After a second filtration at 5 µm with the use of a polypropylene filter cartridge, the water was supplied to the aquariums by a drip irrigation system, the excess being discharged by an overflow. The complete renewal was carried out in approximatively 72 hours, allowing to break the dynamics of the physico-chemical parameters of the incoming outside seawater. Temperature and salinity were maintained stable at 16°C and 38 ppm respectively. Regardless of the photoperiodic conditions applied, the light intensity during the illumination period was 17±3 µmol m^-2^ s^-1^.

##### Manipulation of the light

For the experiment conducted under controlled light conditions, mussels measuring between 50 and 60 mm in total length were selected and their epibionts removed. A total of 450 individuals were distributed in nine aquaria in order to get 50 individuals per aquarium. The aquaria were covered with occulting curtain to prevent the experiment from external light contamination. The mussels were acclimatised for 10 days under an alternating light and dark regime (L:D 12:12, photophase from 06:00 a.m. to 06:00 p.m.) by means of a white lighting ramps (Aquavie, France) coupled with timers. Light intensity at the bottom of the aquarium was of ∼1000 lux. Food was provided once per photoperiodic cycle, at random hours generated using the RAND function of Microsoft Excel. The food consisted of the phytoplankton *Isochrysis galbana* at a final concentration of 3000 cells.mL^-1^ [23]. Phytoplankton were produced by the Mutualised Aquariology Service of the OOB. After the acclimation period, three light conditions were tested in triplicates: constant light (L:L), constant darkness (D:D) and photoperiodic condition L:D 12:12. The food supply was stopped 48 h before sampling. Sampling took place every 4 hours over a period of 28 hours from the 11 December 2019 at 12:00 p.m. to the 12 December at 04:00 p.m. The edge of the posterior mantle region was sampled from five mussels per condition for molecular analysis and stored at −80°C.

##### Manipulation of food availability

Food availability was tested as a time giver, as previously reported in many organisms, notably the clam *Austrovenus stutchburyi* [19,24]. Four conditions were tested: fed once a day at 04:00 a.m. (1xF), fed twice a day at 04:00 a.m. and 04:00 p.m. (2xF), fed continuously (∞xF) and not fed (∅xF). The food consisted of a mixture of three phytoplanktonic strains (1:1:1) *I. galbana, Tetraselmis sp*. and *Rhodomonas salina*. Strains were provided by the Mutualised Aquariology Service of the OOB. The distribution of the food was achieved with the use of an Auto Dosing Pump (Jebao, China). All fed aquaria received a total of 160 mL of the mixture at a concentration of 4.5×10^6^ cells.mL^-1^ per 24 hours. For this experiment, 160 mussels were distributed among four aquaria. Prior to sampling, the mussels were acclimatised for 10 days in constant darkness (D:D) and the experimental condition of food availability assigned. The food supply was stopped 48 h before sampling. Sampling was conducted over 28 hours, with the mantle of five mussels sampled every four hours (beginning at noon) and flash-frozen.

### 2. Gene expression analysis

#### a. Identification of target genes

To construct the target mRNA sequences for *M. galloprovincialis*, a basic alignment search tool (BLAST on Genbank) was used on the sequence read archives (accession number SRX1240182) using known sequences of closely related species available on Genbank. The contigs obtained were aligned and collapsed into a single sequence using the BioLign v.2.0.9 software [25]. The validity of the sequence was assessed by a BLAST on the nucleotide and transcript sequence databases in GenBank. Finally, maximum likelihood trees [26] were constructed from the amino acid sequences using the JTT model [27] with a gamma distribution as implemented in the Mega X software [28]. The statistical robustness of the relationships was assessed using the bootstrap method with 1000 replicates (Supplementary Figures S1-S5). In 2020, a partially annotated assembled genome of *M. galloprovincialis* was released (CGA_900618805.1, Gerdol et al., 2020) and sequences were blast on it (*i.e*., blastx) for cross-validation. If the translated contig matched a hypothetical protein in the genome, a conserved domain search in the pfam database was performed using the NCBI Conserved Domain Search Service [30].

Seven putative genes forming the core of the molecular clock (*i.e., Bmal1, Clock, Period, Cry2, Timeout, Rorb* and *Rev-erb*) were targeted based on the putative molecular clock proposed in the oyster *M. gigas* [12]. Four genes related to biological clocks were also investigated. Cryptochrome 1, CRY1, is involved in the light entrainment of the clock [31,32]. Rhodopsin is a photoreceptor known to have a role in the circadian entrainment of biological clocks, notably in the fruit fly, *Drosophila melanogaster* [33]. Rhythmic expression of *Rhodopsin* has been observed in the mantle of the Arctic scallop *Chlamys islandica* [34]. The third gene is the *Arylalkylamine-N-acetyltransferase, Aanat*, which encodes an enzyme involved in the rhythmic production of melatonin in vertebrates and which secretion and release are rhythmically controlled by biological clocks [35]. A similar observation has recently been made in the labial palps of the razor clam, *Sinonovacula constricta* [36]. The last gene targeted was *HSP70*, which is thought to play a role in the folding and chaperoning of shell matrix proteins [37]. In bivalves, *HSP70* expression varies with temperature [38,39].

Moreover, eight genes related to the biomineralisation process were targeted in aquaria specimens based on the genes used in Louis et al. [6]. Briefly, the targeted genes covered the biomineralisation process at different levels: the transport of mineral components (*i.e., Plasma membrane calcium ATPase* and *Carbonic anhydrase*), the synthesis of the organic matrix (*i.e., Chitin synthase, Chitinase, Tyrosinase* [40,41]), putative inhibitors of mineral matrix formation (*i.e., Perlwapin* and *Nacrein* [42,43]) and a potential regulator of the process (*i.e., Bone morphogenic protein-2 Bmp2* [44,45]).

#### b. Localisation of gene expression by in situ hybridisation

The localisation of targeted gene transcripts in the mussel mantle was realised in individuals sampled at sea in April 2020 at 10:00 a.m. and sacrificed at noon. The posterior edge of the mantle was dissected and fixed in 4% paraformaldehyde (PFA) in phosphate buffer saline (PBS) overnight at 4°C. Tissues were dehydrated in a graded ethanol series (70, 95, 100%), quickly immersed in toluene and then embedded in Paraplast^®^ (Merck, Darmstadt, Germany). Eight-micron thick cross sections (using a MicroM HM 340^E^ microtome, Thermo Fisher Scientific, Waltham, MA, USA) were deposited on glass slides (coated with a 2 % solution of 3−aminopropyl-triethoxy-silane). Prior to *in situ* hybridisation (ISH), sections were successively deparaffinised in toluene, rehydrated (through a series of descending ethanol grades) and placed in PBS.

Five genes were targeted, two related to the biological clock: *Period* and *Clock* and three genes related to the biomineralisation process: *Carbonic anhydrase, Chitinase* and *Bone morphogenic protein-2* (*Bmp2*) using primers described in [6] (Supplementary Table S1). The sequences were subcloned into a pGEX4T1 expression plasmid (Novagen; EMD Chemicals Inc, PA, USA). Anti-sense (AS) and sense (S) riboprobes were produced using the commercial Roche-Diagnostics DIG labelling kit (Merck, Darmstadt, Germany) according to the manufacturer’s recommendations.

The ISH was performed on proteinase K treated sections using the digoxigenin (DIG) labelled antisense and sense probes, as described elsewhere [46]. Briefly, sections were hybridised overnight at 65°C in a hybridisation buffer containing the probes (1µg.mL^-1^). The sections were then washed and incubated overnight in a solution (1/5000) of the anti-DIG antibody conjugated to alkaline phosphatase (Roche, Merck, Darmstadt, Germany). Riboprobes localisations were revealed with Alkaline phosphatase (AP) activity which precipitate the AP substrate (Roche, Merck, Darmstadt, Germany). Sections were observed using an AxioPlan 2 Imaging microscope equipped with a ProgRes® CF^cool^ camera.

#### c. RNA extraction

RNA was extracted using the Maxwell 16 device following the manufacturer’s Simply RNA tissue extraction protocol (Promega Corporation, Madison, WI, USA) for rearing samples reared under controlled light. Due to the low yield of RNA obtained using this technique with mussels, TRIzol-chloroform (Invitrogen, Waltham, MA, USA) extraction was used for all the other experiments.

Briefly, tissues were homogenised in 500 µL of TRIzol using the FastPrep-24 5G (MP Biomedicals, Irvine, CA, USA). The lysate was centrifuged at 10000 rpm for 10 minutes at 4°C. The supernatant was collected and 100 µL of CHCl_3_ was added. The tubes were mixed well before centrifugation at 13000 rpm for 15 minutes at 4°C. The supernatant was kept and 250 µL of isopropanol was added. Samples were then vortexed and incubated for 10 minutes at room temperature before centrifugation at 13000 rpm for 20 minutes at 4°C. The supernatant was discarded and the remaining pellet was washed in 50 µL of 75% EtOH followed a centrifugation at 13000 rpm for 5 minutes at 4°C. This operation was repeated once. The pellet was then air dried at room temperature for 10 minutes before being resuspended in 30 µL of water. DNase treatment was performed using the DNA-free™ kit (Ambion; Austin, TX, USA), following the manufacturer’s protocol.

Concentration and purity were assessed using a NanoDrop spectrophotometer (Nanodrop Technologies, Wilmington, DE, USA) and an Agilent 2100 bioanalyzer (Agilent Technologies, Santa Clara, CA, USA). Gene expression was assessed using the NanoString nCounter™ Gene Expression Assay (NanoString Technologies, Seattle, WA, USA) at the Pôle Technologique CRCT (Toulouse, FR). Probes A and B, between 70 and 90 bp in length, were designed to hybridise to the corresponding target mRNA sequence. The probes set was designed by IDT (Integrated DNA Technologies, Coralville, IA, USA). The total amount of mRNA counted for each gene was normalised following the manufacturer’s guidelines [47].

### 3. Valvometry

Seven mussels were placed in three aquaria (*i.e*., two to three individual/aquarium) for behavioural measurements using valvometry [48,49]. After two days of acclimation under L:D 12:12 condition, the mussels were kept in constant darkness for 12 days, then food (*i.e., I. galbana*) was provided each day at a random time, for eight days. Finally, the same mussels were kept in L:D 12:12 condition with light from 06:00 a.m. to 06:00 p.m. for 13 days. The mussel behaviour was recorded by equipping each specimen with both a magnet and a coated Hall element captor glued to each valve of the shell [49]. For the second part of the experiment (L:D 12:12), one mussel lost its captor. During the experiment, each aquarium was also equipped with a light sensor and a temperature probe for remote condition monitoring. The electrical potential difference measured every second was then converted into a valve opening amplitude (VOA in %) per hour. Valve opening duration (VOD) was determined by calculating the time during which the valves of the mussels were open by more than 20% of the VOA [50,51].

### 4. Chronobiological analysis

Rhythmicity of genes expression was assessed using the R packages “DiscoRhythm” and “RAIN” [52,53] in R (v. 4.1.2) [54]. Cosinor adjustment was realised in DiscoRhythm, allowing to establish acrophase (ϕ) and amplitude (A) data. The RAIN package was used as a complementary non-parametric method designed for biological data integrating different peak shapes [53]. The circatidal range was defined as 12±2 hours and the circadian range as 24±4 hours. Multiple testing deviations were applied using Benjamini-Hochberg corrected p-value at 0.05 [55].

For behavioural analysis, actograms were made using the ImageJ plugin ActogramJ [56]. Prior to rhythm analysis, the absence of randomness in the data and the absence of stationarity were assessed using an autocorrelation function (ACF) and a partial autocorrelation function (PACF) as implemented in R [57,58]. The Lomb-Scargle periodogram was done using ActogramJ with a threshold p-value of 0.05. Significant peaks were validated using a cosinor model as implemented in the “cosinor” and “card” packages in R [59,60]. The model was statistically validated using a goodness-of-fit test, the normality of the residuals and the homogeneity of their variance [58,61]. Rhythm was assessed using an error ellipse test. To assess the presence of an imbricated rhythm, the residuals were reinjected into a cosinor model and the same tests were done. Cross-validation of significant rhythms was done using the R package “RAIN” [53]. Calculations were done at individual and group levels. Rhythmic valve activity was classified according to its period range: circadian, circatidal, bimodal (*i.e*., both circadian and circatidal) or other.

## Results

### 1. Localisation of biological clock and biomineralisation transcripts in the mantle

Using ISH, *Clock* mRNAs were widely detected in the edge of the mantle, *i.e*., in the outer lobe and the basal bulb, in the connective tissue of the inner lobe and in the calcifying epithelium (Figure 1A). In contrast, *period* transcripts were found only in the inner part of the outer lobe (OL), the basal bulb and in the epithelium along the middle lobe. Biomineralisation related genes were expressed in the same area of the mantle as biological clock genes at noon. *Bmp2* transcripts were present at the edge of the basal bulb and in the connective tissue of the inner lobe (IL) (Figure 1B). *Carbonic anhydrase* (*CA*) expression was detected in the basal bulb and along the calcifying epithelium and the inner part of the OL. *Chitinase* gene was expressed in the connective tissue of the IL and a discrete signal was observed in the inner part of the OL. On histological sections, no signal was detected with the sense probes used as negative controls (Supplementary Figure S6).

**Figure 1:**
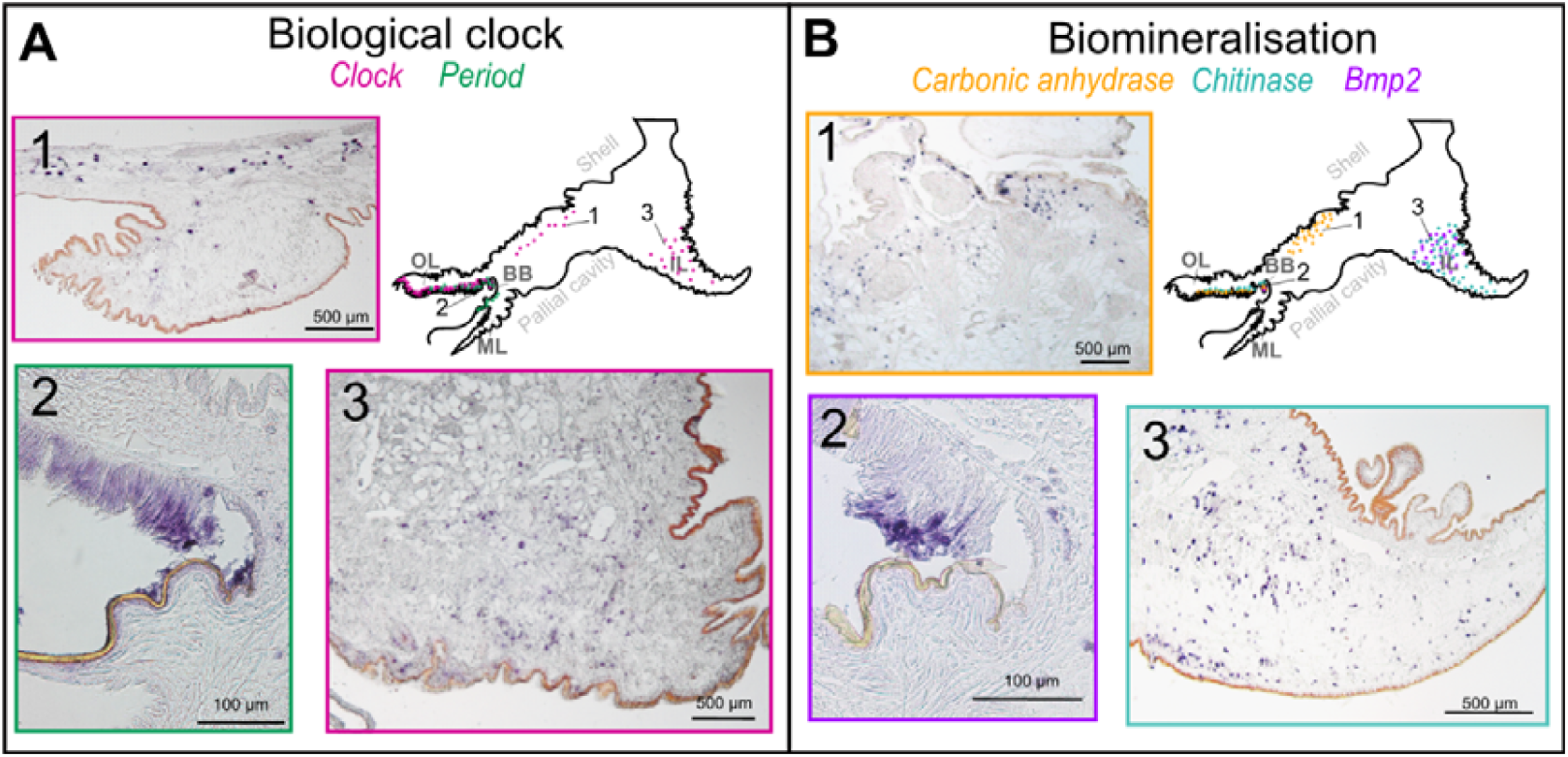
Localisation of the transcripts of the core circadian clock and biomineralisation-related genes in the posterior edge of the mantle of *Mytilus galloprovincialis* revealed by *in situ* hybridisation. A) *Clock* was expressed in the calcifying epithelium (1), in the outer lobe (OL) and in the connective tissue of the inner lobe (IL) (3). *Period* was highly expressed in the basal bulb (BB) (2) and to a lesser extent in the inner part of the middle lobe (ML). B) The three target genes were expressed in the inner part of the OL. *Carbonic anhydrase* was also expressed in the calcifying epithelium (1) and in the BB. *Chitinase* was also expressed in the connective tissue of the IL (3). *Bmp2* was also expressed in the BB (2).

### 2. Gene expression rhythm in *M. galloprovincialis* in the bey of Banyuls-sur-Mer

During the 36 hours of sampling in the bay of Banyuls-sur-Mer, the sun rose up at 07:20 a.m. and set down at 08:10 p.m., solar intensity peaking at 01:44 p.m. The study site at sea was located in an area characterised by a mixed semi-diurnal tidal regime, *i.e*., unequal tidal amplitude with a large amplitude followed by a smaller one. During the sampling period, high tides occurred at 01:55 p.m., 01:14 a.m. and 03:19 p.m.

Among the core biological clock genes, the positive element of the TTFL, *Clock*, showed a significant daily oscillation of expression (RAIN, τ =20h, p=5.2e-5) (Figure 2, Supplementary Table S2), which decreased after midnight and increased after midday (Cosinor, ϕ=12:25 a.m., p=0.001). The expression of *Bmal1*, the second positive element of the loop, was not significantly rhythmic. The expression of *Period*, one TTFL repressor, increased at night and decreased during the day (Cosinor, ϕ=07:25 a.m., p=1e-24), exhibiting a daily expression pattern (RAIN, τ =24h, p=2e-31) with a larger amplitude than other genes that constitute the TTFL (Cosinor, A= 0.93, p=1e-24). The tidal pattern for *Period* expression was significant (RAIN, τ =12 h, p=0.005), although of smaller amplitude than the daily one (Cosinor, A= 0.27, p=0.032), it was characterised by a first peak of expression at 12:35 p.m. (Cosinor, p= 0.032). This gene therefore showed a bimodal rhythm of expression (Figure 2). In contrast, CRY2, which presumably forms a trimer with PERIOD and TIMEOUT proteins, showed no significant rhythmic expression. Gene expression levels for *Timeout* and *Rorb* could not be measured because mRNA levels were below the limit of detection.

**Figure 2:**
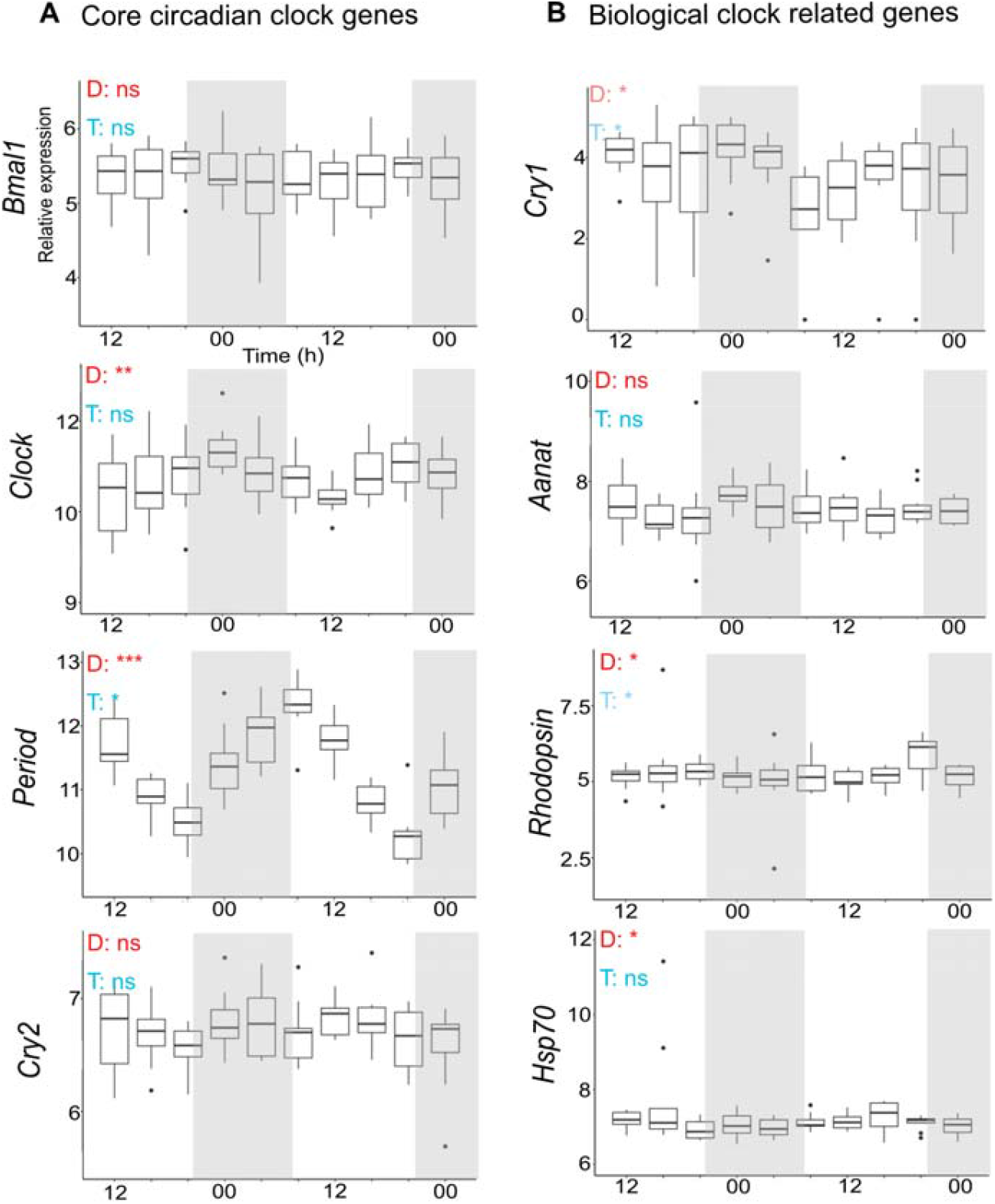
Core circadian clock and related gene expression at sea (SOLA). Rhythmicity of expression was tested for daily range of expression (D) and tidal range (T). Cosinor adjustment and RAIN analysis were performed to test rhythmicity of expression. If only one was significant, the significance level is shown in a lighter colour. Grey areas indicate nighttime and white ones indicate daytime. Significance levels: ns: p>0.05; *: 0.05 ≥ p ≥ 0.01; **: 0.01 > p ≥ 0.001; ***: p <0.001

Looking at biological clock related genes, *Cry1*, which has a putative role in light entrainment of the clock, showed a large variability between replicates compared to other clock related genes (Figure 2). A discrepancy between the adjustment models was observed for this gene: the RAIN adjustment showed a significant daily gene expression (RAIN, τ =28 h, p=0.028) while the cosinor adjustment showed a tidal expression (Cosinor, τ =14h, p=0.020). The photopigment gene *Rhodopsin* showed a daily expression (RAIN, τ =24h, p=0.018) with an acrophase at 05:35 p.m. (Cosinor, p=0.042). The significant tidal component observed for this gene using the cosinor adjustment had a close similar acrophase (Cosinor, ϕ =06:00 p.m., p=0.05) and was less significant (Cosinor, τ =13h, p=0.05). *Aanat* showed no significant rhythmic expression. *Hsp70* had a daily expression (RAIN, τ =28h, p=0.008) with a relatively small amplitude (Cosinor, A=0.17, p= 0.038) and a peak of expression at 03:15 p.m..

### 3. Gene expression in *M. galloprovincialis* reared in controlled conditions

#### a. Gene expression in mussels reared under controlled light conditions

Mussels were reared under three lighting conditions: alternation of 12 h of light and 12 h of darkness (L:D 12:12), constant darkness (D:D) and constant light (L:L). Under photoperiodic condition L:D 12:12, among the core biological clock genes, *Clock* (RAIN, τ =24h, p= 0.042) and *Period* (Cosinor, τ =20h, p=0.047) showed significant daily oscillations in their expression, which profiles were antiphasic with a peak of expression reached at 03:50 p.m. for the activator *Clock* (Cosinor, p>0.05) and at 04:25 a.m. for the repressor *Period* (Cosinor, p=0.047). None of the other targeted genes showed significant rhythmicity of expression in L:D 12:12 condition. Different responses were observed between the two constant light conditions. In L:L, all core biological clock genes showed significant daily cyclic expression except *Bmal1. Clock* (Cosinor, ϕ=12:35 p.m., p = 3.9e-4) and *Period* (Cosinor, ϕ=01:45 p.m., p = 0.43) exhibited an expression profile with a similar acrophase around noon. The second repressor, *Cry2*, had a peak of expression at 09:30 p.m. (Cosinor, p>0.05) (Supplementary Figure S8). Among the biological clock related genes, *Rhodopsin* also showed circadian rhythmic expression (RAIN τ =28h, p=0.009), whereas *Hsp70* had circatidal expression (Cosinor, τ =11h, p=0.022) (Supplementary Table S4). In D:D, only the two positive elements of the biological clock, *Clock* and *Bmal1*, showed significant oscillations in expression. *Bmal1* showed a circadian expression pattern (Cosinor, τ =23h, p=0.011) with an acrophase at 12:45 p.m. and *Clock* had a circatidal expression (Cosinor, τ =14h, p=0.012), with an acrophase at 06:00 p.m, and a less significant circadian expression, with an acrophase at 02:10 p.m. *Hsp70* showed also a bimodal expression with circadian (RAIN, τ =28h, p=0.036) and circatidal (Cosinor, τ =14h, p=0.042) components, both with an acrophase at 03:35 p.m. (Supplementary Table S5).

No rhythmic expression was detected for all the biomineralisation related genes studied under L:D 12:12. However, under constant light condition (L:L or D:D), both circadian and circatidal oscillations in gene expression were observed (Figure 3; Supplementary Table S3). In both conditions, *Carbonic anhydrase* and *Nacrein* had circadian expression with a period of 24 hours in L:L and 28 hours in D:D. *Chitinase* had a circatidal expression in both conditions with a period of 14 hours in L:L (Cosinor, p=0.015) and 10 hours in D:D (Cosinor, p=0.036). Another gene involved in chitin metabolism, *Tyrosinase*, had a significant circatidal expression in D:D (Cosinor, τ =14h, p=0.001). A significant circatidal oscillation was also observed for *Ca*^2+^ *ATPase* in the L:L condition (Cosinor, τ =10h, p=0.015).

**Figure 3:**
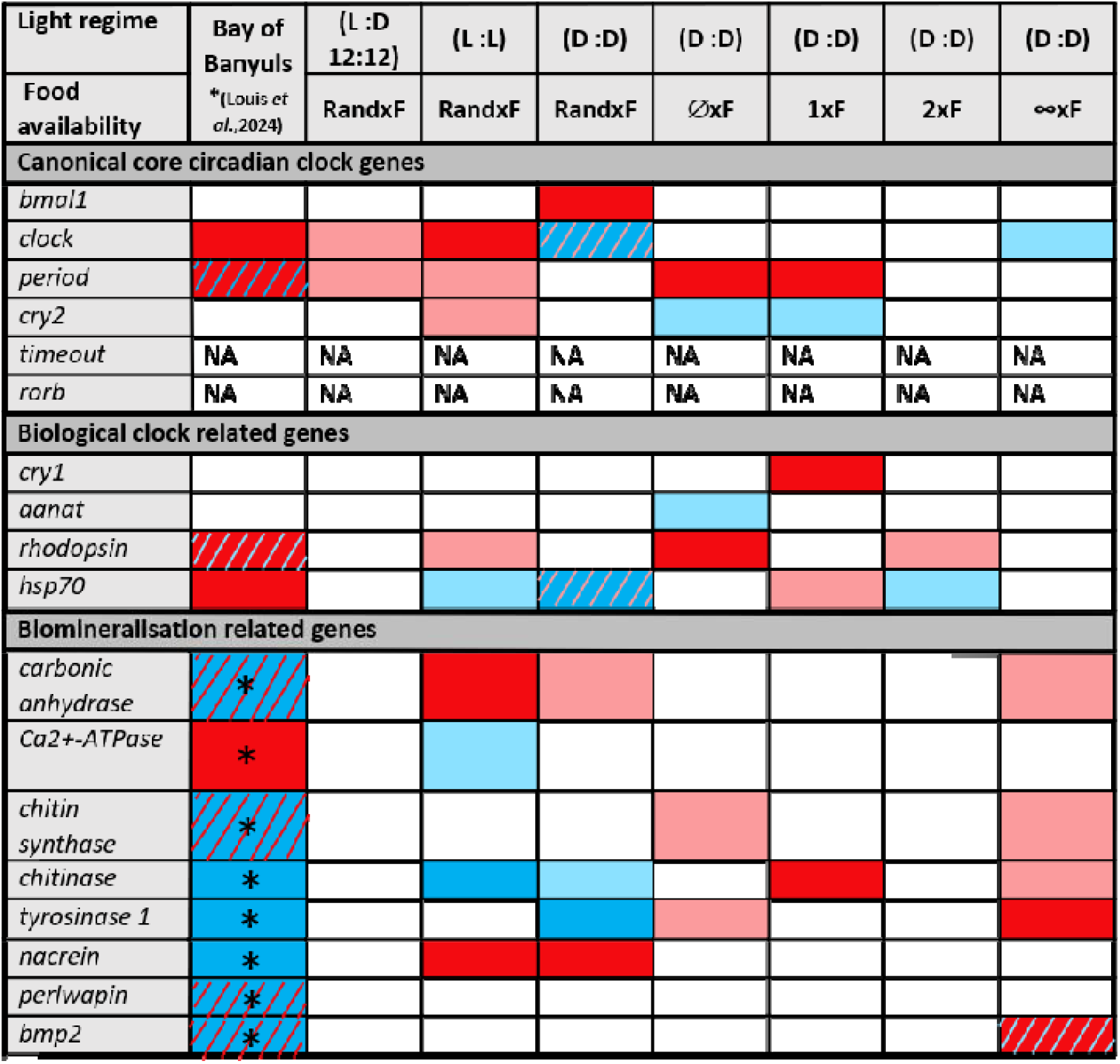
Rhythmicity of expression of genes related to biomineralisation and biological clocks under manipulated conditions of light and food availability. Circatidal rhythmicity is shown in blue and circadian rhythmicity is shown in red. Two mathematical adjustments (Cosinor and RAIN) were used to test for rhythmicity in gene expression. When one of the two adjustment was significant, light blue or light red are used. Bimodal rhythmicity is shown in two-colour texture, with the predominant colour corresponding to the main rhythm. When no significant rhythmicity was observed, the case is in white. No available data are annotated “NA”. Measurements taken in the Bay of Banyuls-sur-Mer are marked with an asterisk (*) and were retrieved from Louis *et al*., 2024. Three light conditions were tested; continuous darkness (D:D), continuous light (L:L) and alternation of light (12 h) and dark (12 h) (L:D 12:12) crossed to a random distribution of the food (RandxF). Four conditions of food availability conditions were crossed with D:D condition: fed once a day (1xF), fed twice a day (2xF), fed continuously (∞xF) and never fed (∅xF).

#### b. Gene expression in mussels reared under different food availability conditions

The food availability cue was tested as time giver using four feeding conditions (fed one a day (1xF), twice a day (2xF), continuously (∞xF) or never fed (∅xF). Among the biological clock related genes in the 1xF condition, *Cry1* (Cosinor, τ =20h, p=0.012) and *Hsp70* (Cosinor, τ =28h, p=0.030) had daily expression (Supplementary Figure S11). In the 2xF condition, *Hsp70* had significant tidal expression (Cosinor, τ =12h, p=0.047) and *Rhodopsin* had circadian expression (RAIN, τ =28h, p=0.038). When not fed, *Rhodopsin* also had a significant circadian expression (RAIN, τ =20h, p=0.020), which was not the case in the other constant condition. *Aanat* was observed as a circatidal expression in the ∅xF condition (Cosinor, τ =10h, p=0.027).

Rhythmicity was lost for most biomineralisation genes when fed once a day (1xF), twice a day (2xF) and when never fed (∅xF) compared to the marine environment (Figure 3) [6]. For the mussels in 1xF condition, only *Chitinase* was expressed according to a circadian pattern (RAIN, τ =20h, p=0.008) with a peak at 04:05 a.m., which corresponds to the feeding time (Cosinor, p=0.047) (Supplementary Figure S12). For mussels in 2xF condition, none of the genes showed significant oscillation in expression over time. In both constant feeding conditions, more rhythmic genes were observed. Under ∅xF condition, only the genes *Tyrosinase* (RAIN, τ =28h, p=0.004) and *Chitin synthase* (RAIN, τ =20h, p=0.016) showed a significant circadian periodicity of expression. When fed continuously (∞xF), most of the biomineralisation related genes had a significant circadian rhythm of expression (*i.e., Carbonic anhydrase, Bmp2, Chitin synthase, Chitinase* and *Tyrosinase*, Figure 3).

Looking at core biological clock genes, similar results were observed when in 1xF and ∅xF conditions (Figure 3; Supplementary Tables S6-S9). In both conditions, *Period* showed an oscillation of expression with a period of around 24 hours and *Cry2* showed a circatidal expression (Supplementary Figure S10). Under 2xF condition, none of the core clock genes showed significant oscillations in expression. Whereas under ∞xF condition, *Clock* had circatidal expression (Cosinor, τ =12h, p=0.033).

### 4. Valve aperture analysis in mussels reared under controlled light conditions

Valve activity was recorded for a group of mussels (N=7) sampled in the bay of Banyuls at 9 a.m.. Following their capture, they were immediately placed in an aquarium under constant darkness (D:D) for 12 days and then switched to photoperiodic condition (L:D 12:12) (Figures 4). In D:D condition, the rhythm of activity had a smaller amplitude than in L:D and was bimodal at the group level (RAIN, τ =12h, p=1.46e-09 and τ = 27h, p=9.35e-10). Five out of seven mussels showed no rhythmic behaviour. Of the two rhythmic individuals, one showed a circadian behaviour with a period of 26 hours and the other one a circatidal behaviour with a period of 12 hours. Under photoperiodic conditions (L:D 12:12) the mussels showed a daily activity at the group scale, their valves being closed during the day and open at night (RAIN, τ =24h, p=3.52e-57). At the individual scale, the majority of the mussels (4/6) showed a daily behaviour with, however, different acrophase times according to each rhythmic animal, suggesting they were not synchronised among the sampled population. Finally, one other mussel showed a circatidal behaviour and the last one exhibited no behavioural rhythm. Interestingly, the mussel that showed circatidal behaviour was the same individual in both conditions.

**Figure 4:**
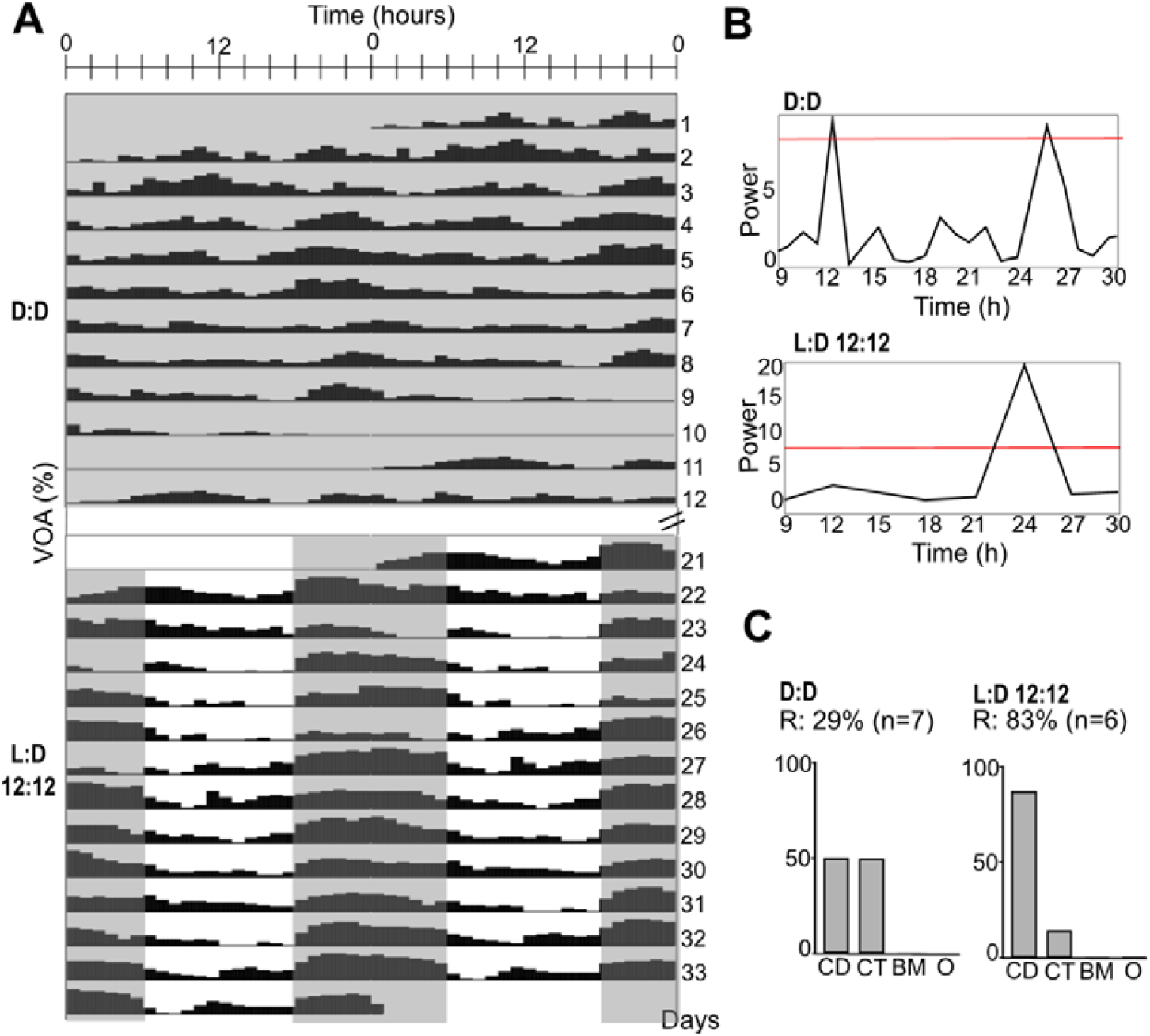
Valve activity of mussels reared under two successive lighting conditions. A) Double plotted actogram of mean valve opening amplitude (VOA) of mussels (N = 7) reared under two photoperiodic conditions. Valve opening amplitude was averaged in percent per hour. Mussels were first kept in constant darkness (D:D) for 12 days. After 8 days of random feeding in D:D, they were placed under light-dark alternation condition (L:D 12:12, alternation of 12 hours of light and 12 hours of darkness) for 13 days. Grey shading corresponds to the dark phase. B) Lomb-Scargle periodograms constructed with the mean values of VOA for each photoperiodic condition, peaks above the red line are significant. C) Behavioural rhythmicity expressed at the individual level, rhythmicity was tested and classified for rhythmic individuals (R, in percent) in circadian (CD), circatidal (CT), bimodal (BM) or other (O) periods.

## Discussion

### 1. *Mytilus galloprovincialis* biological clocks sustain circadian and circatidal rhythms

Before assessing the relation between biomineralisation and biological clock it is necessary to determine if *M. galloprovincialis* has a functional clock entraining circadian and circatidal rhythms. We identify the core activators *Clock* and *Bmal1* and the core repressors *Period, Cry2* and *Tim* in *M. galloprovincialis* as reported in other bivalves [12,34,62–64]. *Rorb* and *Rev-erb* may form a second feedback loop, among which only the *Rorb* sequence could be identified. In specimens sampled at sea, both the positive element *Clock* and the negative element *Period* are expressed according to a daily pattern, probably synchronised with the photoperiod as it is known to be the main time giver of the circadian clock [18]. *Period* also exhibits a tidal pattern of expression, characterised by a peak around high tide, and a smaller amplitude than the daily oscillation. Interestingly, *Cry1* which is known as a blue light sensor in Monarch butterfly and to mediate the degradation of the repressor TIM, shows a bimodal expression variation in *M. galloprovincialis* [65].

To characterise the biological clock in *M. galloprovincialis*, experiments were performed under controlled environmental conditions using the photoperiod and the food availability as time giver. When entrained with daily cues (*i.e*., LD 12:12 or 1xF), rhythmic circadian core genes are mainly expressed with daily oscillations. Similar results are observed in the experiment on valve activity for mussels exposed to daily oscillations. However, under tidal cues (*i.e*., 2xF), no significant periodic oscillation of circadian core gene expression is detected. It is possible that some other tidal environmental variables, such as currents or water pressure, are stronger tidal time givers and can counterbalance the entrainment by the light [66]. Under constant light condition (*i.e*., L:L), most of the core clock genes exhibit circadian oscillations which was not the case in constant darkness (*i.e*., D:D). Similarly, stronger circadian activity was observed under constant light than in constant darkness in the amphipod *Parhyale hawaiensis* [14]. In other free-running conditions (*i.e*., D:D, ∅xF and ∞xF) both circadian and weaker circatidal expressions are observed. This suggests that the expression of some constituents of the core circadian clock (*i.e., Cry2* and *Clock*) might be plastic and can shift from circadian to circatidal in *M. galloprovincialis*. This corroborates observations made in the Pacific oyster *M. gigas* as in specimens entrained under tidal regime, the core circadian clock gene expression shift from circadian to circatidal expression [16]. However, knock down of *Per, Cry2* and *Clock* using RNA interference in the mangrove cricket *Apteronemobius asahinai* and the isopod *Eurydice pulchra* disrupted circadian outputs but not the circatidal ones [15,67–69]. Up to now, only BMAL1 has been shown to be shared in *P. hawaiensis* and *E. pulchra* [14,15]. Therefore, based on the literature it is unclear if all core circadian clock genes are required for circatidal rhythms in marine organisms.

As observed in numerous organisms, both terrestrial and aquatic, the rhythmic behaviour observed in *M. galloprovincialis* is controlled by biological clocks [70,71]. It is interesting to note that a plasticity similar to that observed at the molecular level was also identified at the behavioural level. Mussels collected at sea and reared in D:D condition show a bimodal pattern of valve activity in the sampled group. Of the two individuals (over seven) in which rhythmic valve activity persists under D:D conditions, one exhibits a circadian behaviour and the other, a circatidal one. This confirms the endogenous nature of both types of rhythm in *M. galloprovincialis* and that this species possesses a functional clock.

In all measurements done, the circatidal rhythm was fainter than the circadian one and this observation could be related to the mussel environment. In the bay of Banyuls-sur-Mer, mussels are exposed to a microtidal tidal regime, associated to a mixed semidiurnal tidal regime. The clock is thus probably poorly entrained to tides, leading the photoperiod as a stronger time giver for clock genes in *M. galloprovincialis* in this area. Repeating these experiments in *M. galloprovincialis* specimens inhabiting locations with a semi-diurnal tidal regime, such as those found along the Atlantic coast, could provide valuable insights into the interplay between the circadian and circatidal rhythms.

### 2. Biological clocks do interact with the biomineralisation process, but the nature of the interaction remains elusive

At the functional level in *M. galloprovincialis*, the present study reveals a co-expression of the core biological clock genes, *Clock* and *Period*, together with three genes related to biomineralisation in the posterior edge of the mantle, known as the organ of biomineralisation in bivalves [1]. Interestingly, all the biomineralisation and biological clock detected transcripts are expressed in areas of the mantle that are involved in the biomineralisation process, such as the basal bulb which secretes the periostracum, or the outer lobe, known for the secretion of shell constituents, or the calcifying epithelium [72,73]. *Clock, Chitinase* and *Bmp2* transcripts are also observed in the inner lobe (IL). As the IL is the first tissue in contact with sea water entering the mantle cavity, it is likely that the presence of transcripts of biomineralisation-related genes such as *Chitinase* in the IL epithelium is related to its role in immunity through hydrolysis of chitin-coated microorganisms [72,74]. However, their expression within the IL may also be related to the haemolymph sinuses present within the mantle tissue, as haemocytes H2 and H3 have been shown to express several genes related to organic shell compounds [43,75–77]. Given the joint presence of transcripts linked to both biological clock and biomineralisation in key places for the formation of the shell, this supports the hypothesis that biological clock may orchestrate the growth of hard tissue in *M. galloprovincialis*.

As previously reported [6], the shell increment patterns and biomineralisation gene expression of *M. galloprovincialis* primarily reveal a growth following tidal rhythms in the bay of Banyuls-sur-Mer. In the present study, core biological clock genes exhibit strong daily variations, that are slightly modulated by weak tidal oscillations occurring in the bay. Therefore, it seems that the biological clock and the biomineralisation process of mussels do not oscillate at the same pace. In marine ecosystems, organisms are exposed to a wide range of rhythmic environmental cues (*i.e*., photoperiod, temperature, tides) so that the output of the endogenous clock can be masked or attenuated by direct responses to environmental variations [78,79]. Consequently, the results we obtained with the mussels sampled in the field must be used with caution and do not allow us, yet, to totally rule out the role of the endogenous biological clock in the control of the rhythmic dynamics of shell biomineralisation in *M. galloprovincialis*.

In mussels reared in controlled environmental conditions at the lab, under both constant photoperiodic conditions (*i.e*., LL and DD) and constant feeding conditions (*i.e*., ∅xF and ∞xF), significant circatidal and circadian rhythmic expression of biomineralisation-related genes is observed, suggesting the endogenous nature of the rhythm. This observation demonstrates that the biomineralisation process is probably linked to endogenous timekeeping mechanisms. This is in line with observations made in *Cerastoderma edule* shell, in which growth pattern are still formed under constant environmental conditions [8]. Curiously, no rhythmic expression of targeted biomineralisation genes followed the LD cycle nor the 2xF cue, which is similar to what is observed in circadian clock genes. This could suggest that the activation of the transcription of biomineralisation genes is controlled by the biological clock. However, discrepancies in the expression patterns of biological clock and biomineralisation genes are observed in a constant environment. For example, when mussels were continuously fed and placed in constant darkness, the expression of the biomineralisation genes as a whole take on circadian expression, whereas the only core biological clock gene cycling, *Clock*, exhibits a rhythmic circatidal expression. Therefore, the interaction between the biological clock genes and the targeted biomineralisation ones is probably not as direct as expected. Different hypothesis can be advanced. First, the core circadian clock genes are not necessary for circatidal rhythm in mussels. This is unlikely as there are growing evidence that circadian clock genes have been co-opted to generate circatidal rhythms in marine organisms such as oysters and crustaceans [14–16,80]. Second, the cells situated in the different area of the mantle (*i.e*., outer lobe, calcifying epithelium and basal bulb) do not have a synchronised expression in mussels reared in aquaria due to missing time giver which are present in their natural environment (*i.e*., temperature, vibration). A closer inspection of expression patterns in the mantle tissue using ISH or single-cell approaches might show different expression rhythms in function of the structure observed. Third, the control of biomineralisation gene expression by the biological clock does not occur directly through gene expression pathways in the mantle, but rather indirectly like in another tissue involved in the synthesis of biomineralisation components [81,82]. The last hypothesis is an indirect effect of the biological clock on the biomineralisation process, involving the motor activity of the mollusc valves. Indeed, it is known that the opening/closing activity of the valves is under the control of the biological clock [12,83]. It is also known that valve closure induces an internal pH drop and subsequent shell decalcification [84]. The biomineralisation pathway could thus be activated or repressed by internal physiological conditions such as internal pH or ion concentration variation in the extrapallial fluid. Further experiments should be carried out to better understand whether biological clocks directly or indirectly drive the rhythm of the biomineralisation process.

In summary, our study suggests that biological clock may control the biomineralisation process as previously suspected based on observation of increment patterns from the shell of *M. galloprovincialis* sampled in different Mediterranean environments [6]. The key environmental factor seemed to be the food availability. However, biomineralisation genes expression of mussels being fed once or twice a day did not reflect the rhythm of food availability. This shows that the interaction between the biological clock and the biomineralisation process is complex and probably integrate many factors that remain to be discovered. It would undoubtedly be appropriate to consider a non-targeted approach, by sequencing the whole transcriptome of *M. galloprovincialis* mantle after an entrainment to daily or tidal cues.

### 3. Implications of the effect of biological clocks when using bivalve shells as biological archive

Shells of actual and fossil bivalves are widely used to reconstruct present and past environment as the observation of shell growth patterns give information on the environmental parameter that are estimated from shell chemical composition. This makes bivalve shell a biological archive. The study of the influence of the biological clock in the biomineralisation process is crucial to understand how it could be a potential source of error for climatic reconstructions. Indeed, biological clock control could influence the shell growth pattern formation, but it also might influence ion uptake during the biomineralisation process, leading to the incorrect estimation of environmental parameters. Discrepancies between growth patterns and the cyclic environment were observed in our previous *in-situ* study [6], pointing out that errors possibly related to non-alignment of the biological clock and direct environmental biomineralisation drivers in environments without strong tidal variations such as the Mediterranean Sea. Importantly, this is also the case in an environment perturbated by anthropogenic activities as it has been shown that artificial light at night (ALAN) can disrupt the circadian clock of intertidal animals [85,86]. Similarly, water vibrations caused by boat traffic or offshore wind farm might affect the circatidal clock even though it has never been studied until now.

In conclusion, such analyses are fundamental for the validation of bivalve shells as biological archives in sclerochronology, to frame ecological approaches of mollusc populations.

## Acknowledgements

We acknowledge the facilities of Biology platform of imaging (BioPIC). We are grateful to the Bio2Mar platform (http://bio2mar.obs-banyuls.fr) for providing access to instrumentation. We acknowledge Michel Groc from Department of Information and Communication Systems at the OOB and his student Lucas Laveissiere (IMERIR) for the development of the valvometry device. We are thankful to Nancy Trouillard and Pascal Romans at the Mutualised Aquariology Service (OOB) for providing aquariology expertise and phytoplankton strains. We are thankful to captain Eric Martinez, and the crew from the oceanographic boat Nereis II as well as the divers, Jean-Claude Roca and Bruno Hesse (Sea Service from OOB) for the sampling effort at sea. We would like to thank Thomas Moura for his help comparing classical qPCR approach with the Nanostring within the framework of his Master 2 internship.

## Supplementary Figures and Tables

**Figure S1:**
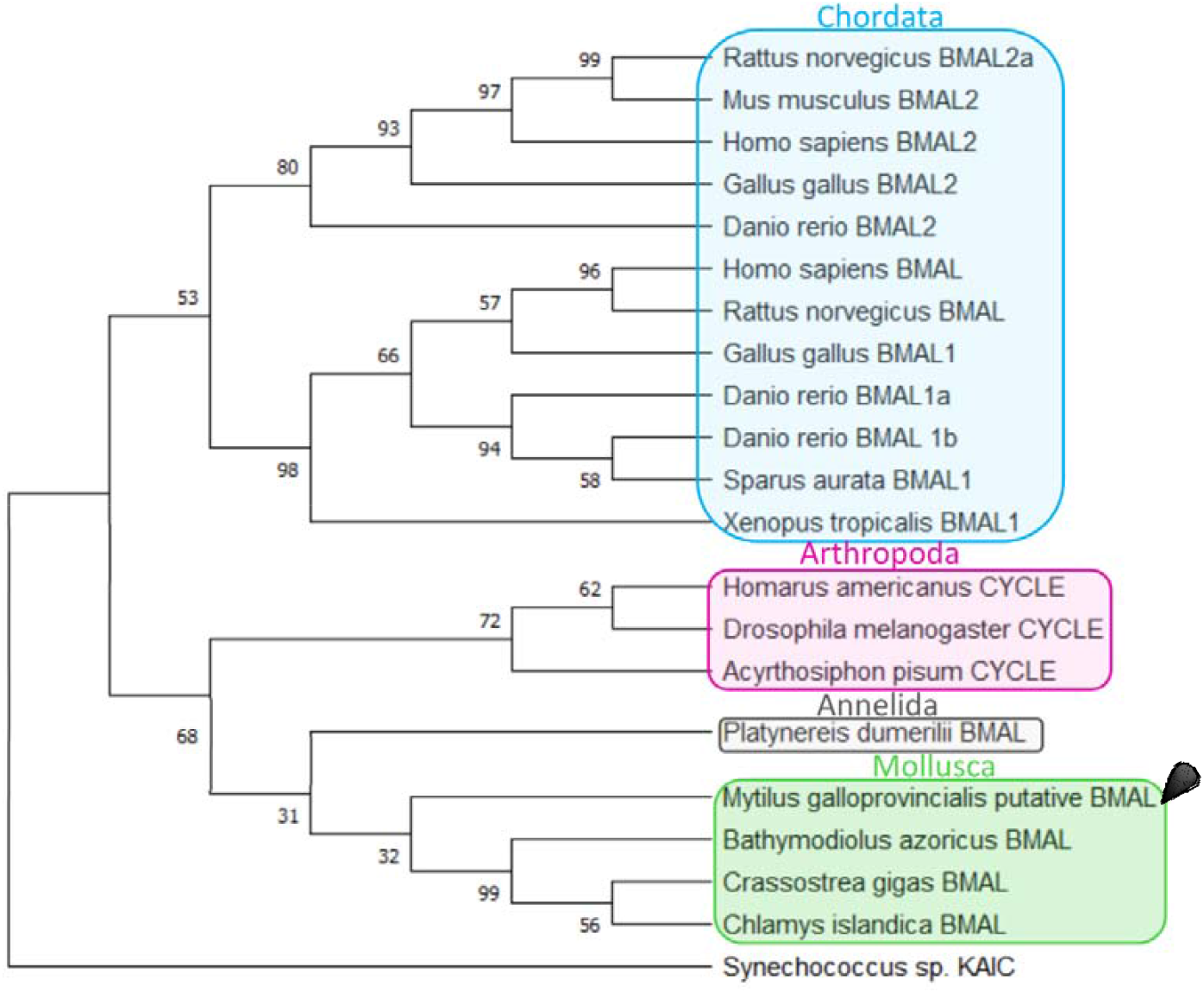
Phylogenetic tree based on BMAL1 sequences. The tree was generated using the maximum likelihood method with the JTT matrix-based model and the gamma distribution. Percentage of bootstraps were based on 1 000 replicates.

**Figure S2:**
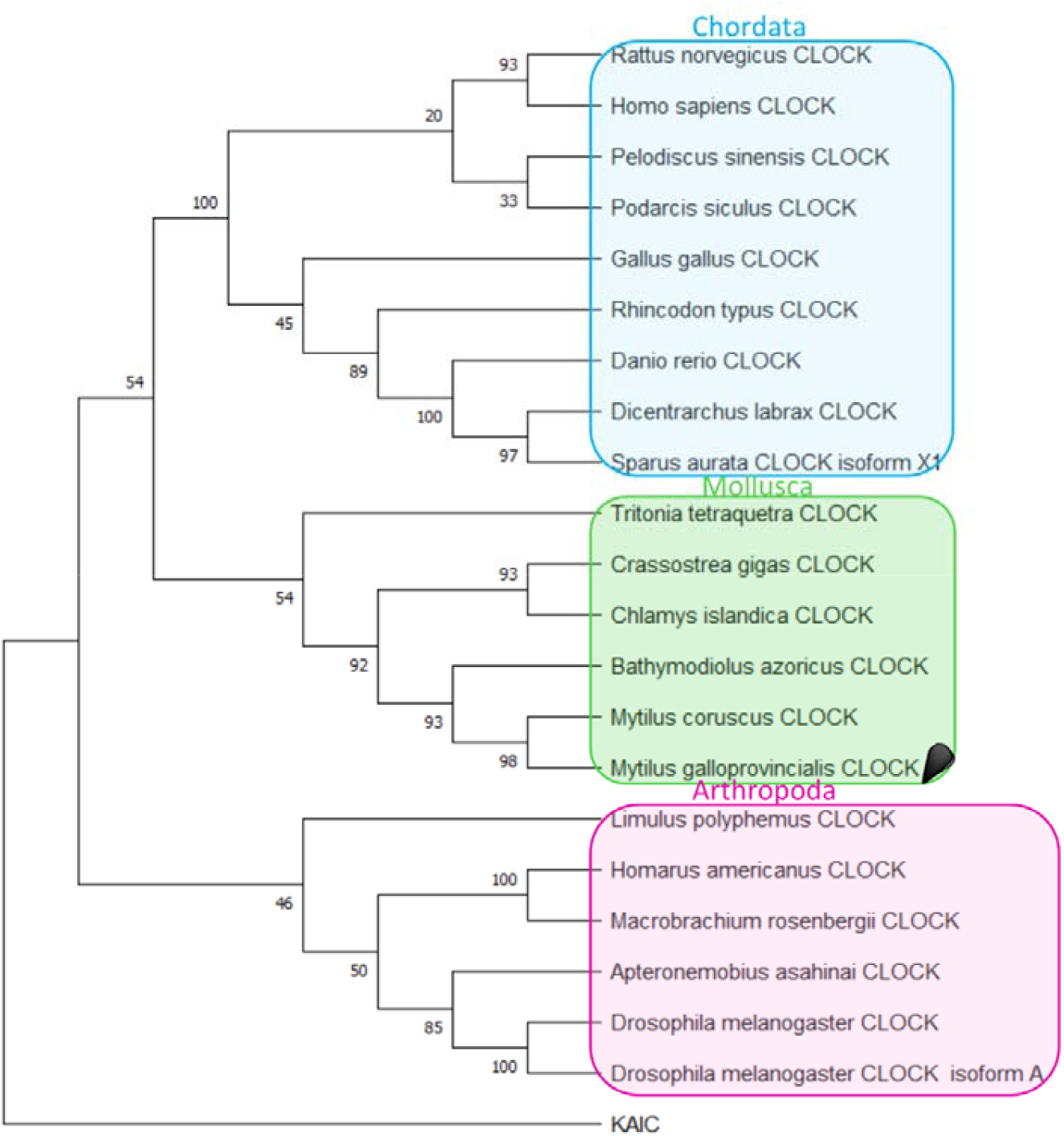
Phylogenetic tree based on CLOCK sequences. The tree was generated using the maximum likelihood method with the JTT matrix-based model and the gamma distribution. Percentage of bootstraps were based on 1 000 replicates.

**Figure S3:**
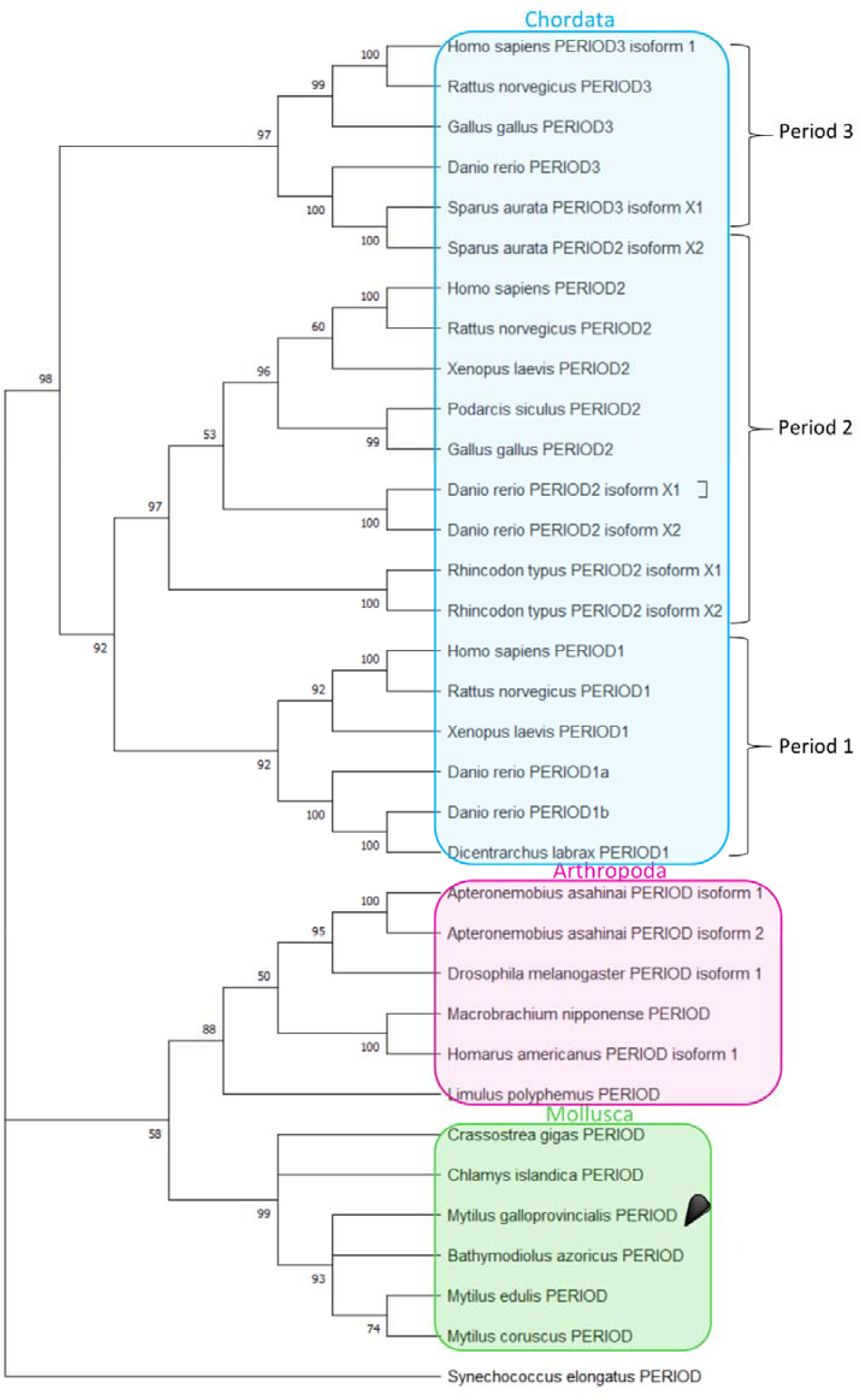
Phylogenetic tree based on PERIOD sequences. The tree was generated using the maximum likelihood method with the JTT matrix-based model and the gamma distribution. Percentage of bootstraps were based on 1 000 replicates.

**Figure S4:**
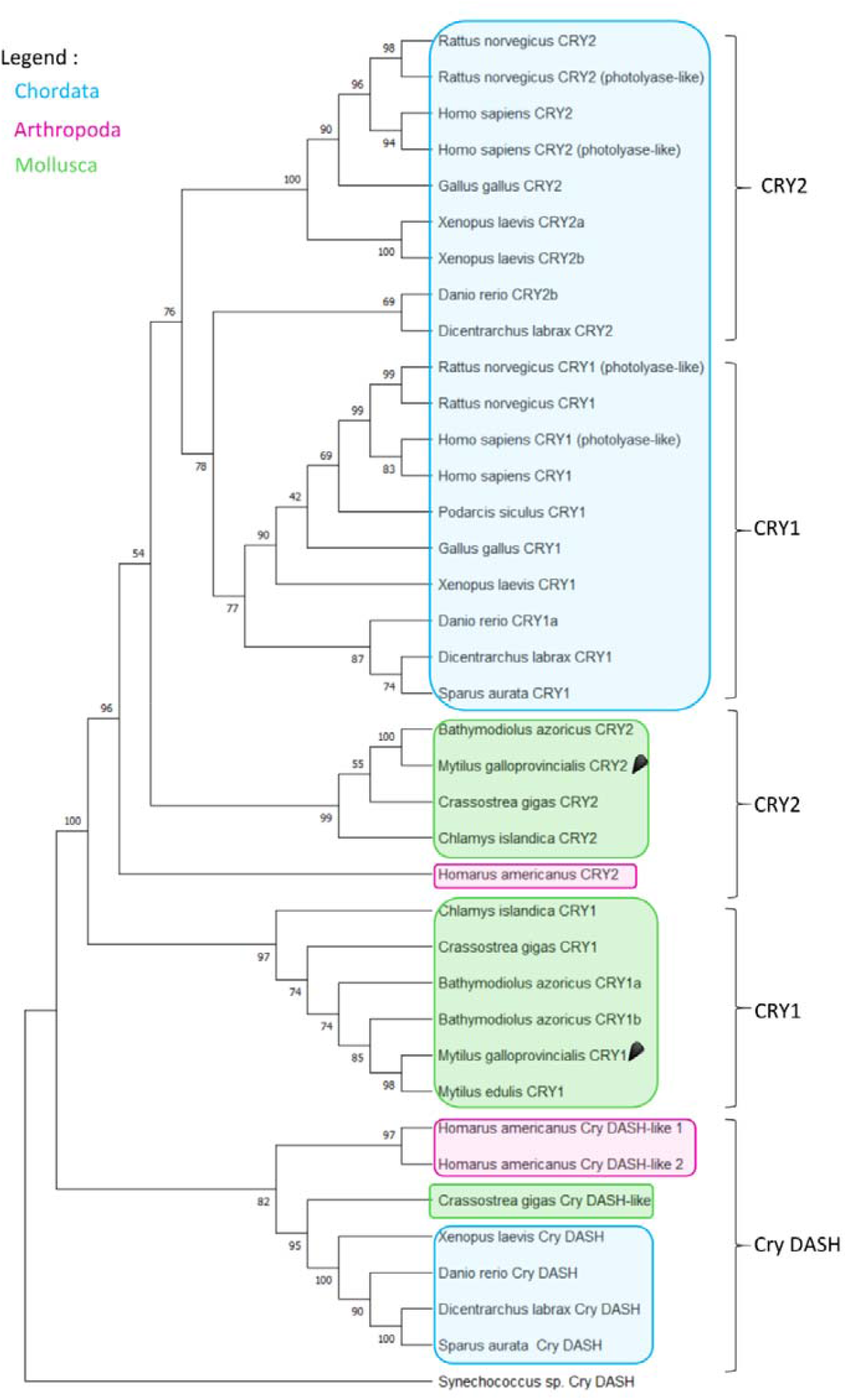
Phylogenetic tree based on CRYPTOCHROME sequences. The tree was generated using the maximum likelihood method with the JTT matrix-based model and the gamma distribution. Percentage of bootstraps were based on 1 000 replicates.

**Figure S5:**
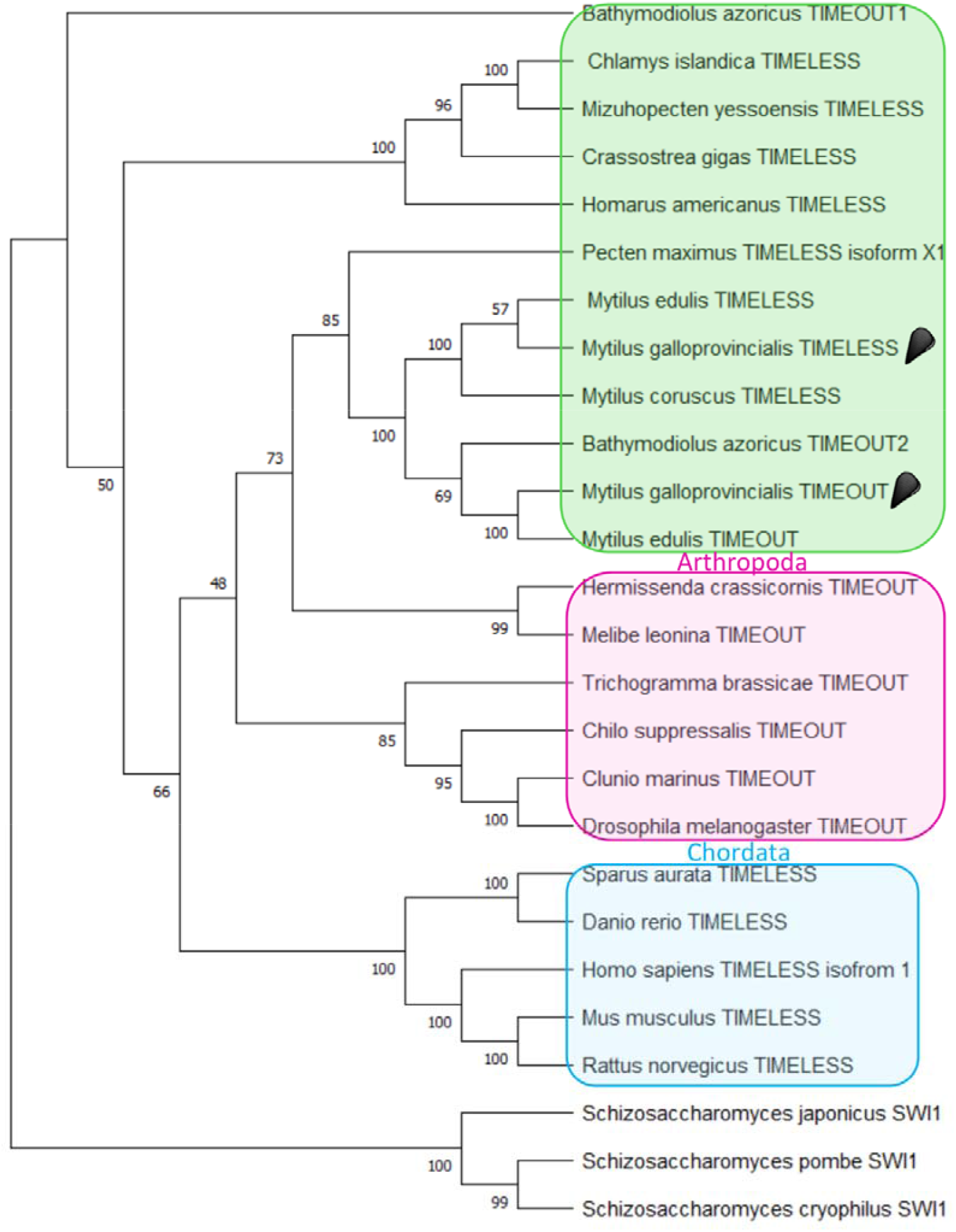
Phylogenetic tree based on TIMEOUT and TIMELESS sequences. The tree was generated using the maximum likelihood method with the JTT matrix-based model and the gamma distribution. Percentage of bootstraps were based on 1 000 replicates.

**Figure S6:**
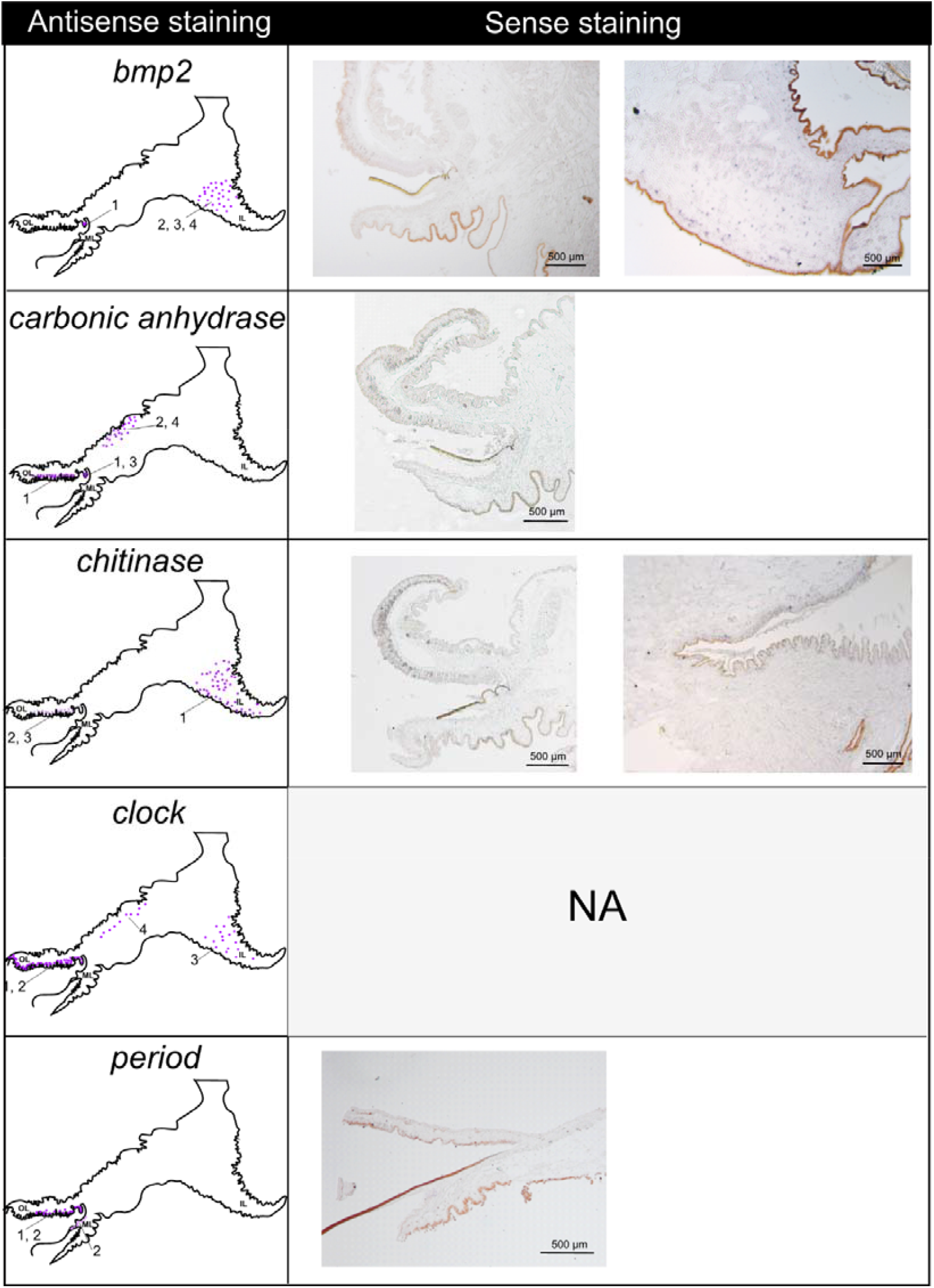
Negative controls of *in situ* hybridisation. Sense probes of *bmp2, carbonic anhydrase, chitinase* and *period* showed no signal. *Clock* sense probe was not specific (ND).

**Figure S7:**
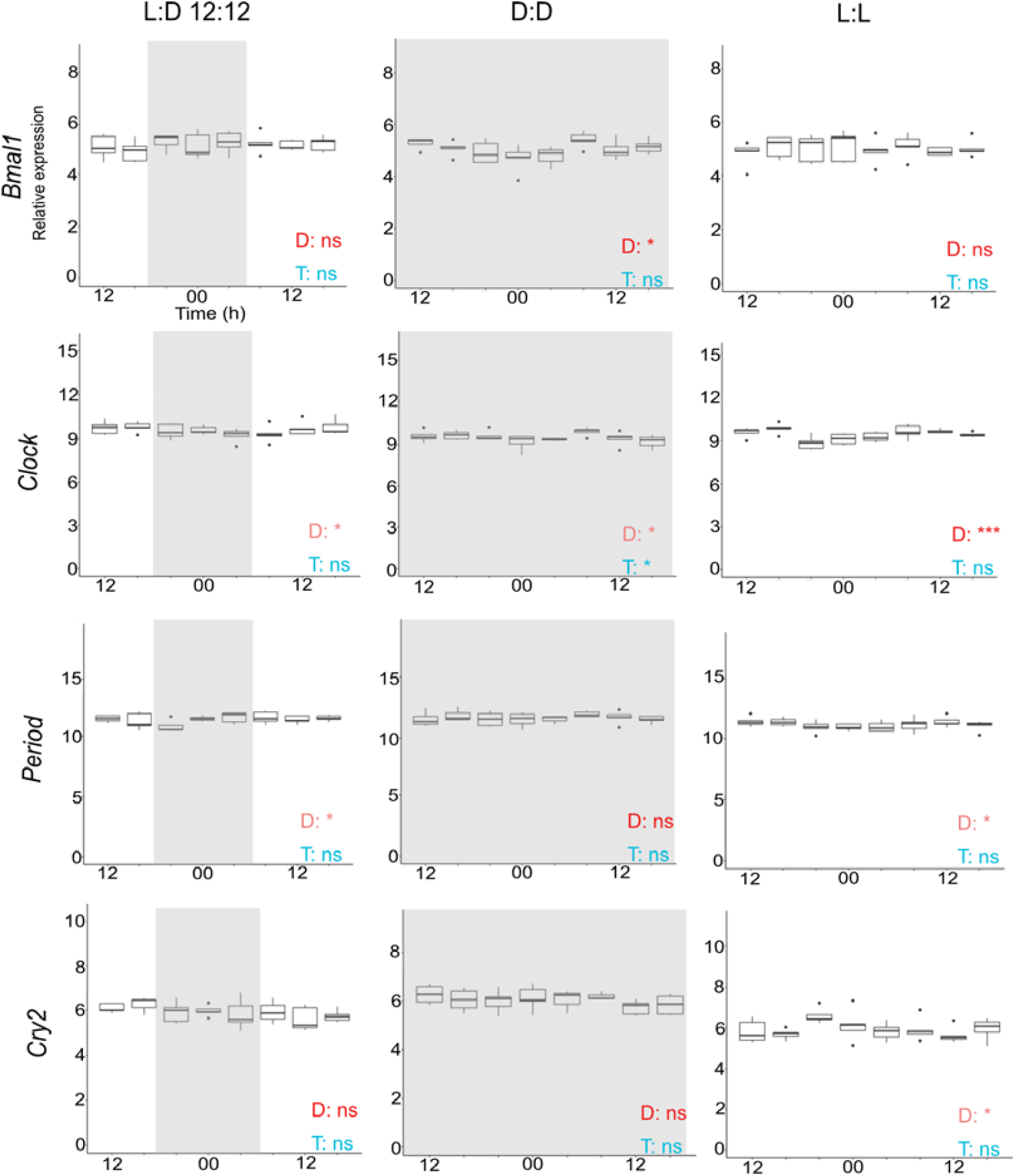
Core circadian clock gene expression under controlled condition of light. Light conditions tested were photoperiodic condition of an alternance of 12 hours of light (in white) and 12 hours of dark (in grey) (L:D 12:12), continuous darkness (D:D) and continuous enlightenment (L:L). Rhythmicity in expression were tested for the daily range of expression (D) and the tidal range (T). Cosinor adjustment and RAIN analysis were achieved to test rhythmicity of expression. When only one was significant, the significance level is in lighter colour. Significance levels: ns: p>0.05; *: 0.05 ≥ p ≥ 0.01; **: 0.01 > p ≥ 0.001; ***: p <0.001

**Figure S8:**
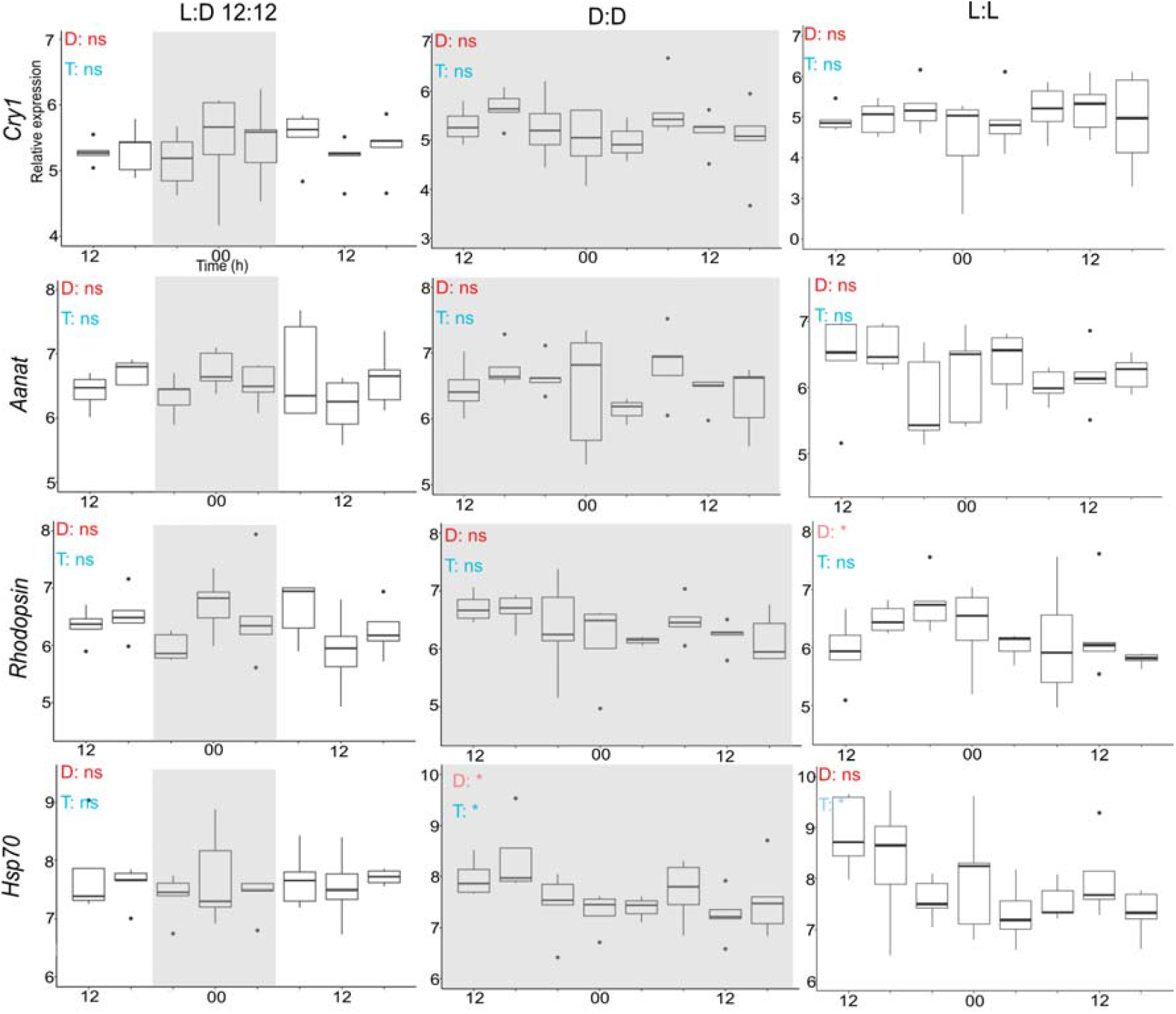
Expression of genes related to biological clock under controlled condition of light. Light conditions tested were photoperiodic condition of an alternance of 12 hours of light (in white) and 12 hours of dark (in grey) (L:D 12:12), continuous darkness (D:D) and continuous enlightenment (L:L). Rhythmicity in expression were tested for the daily range of expression (D) and the tidal range (T). Cosinor adjustment and RAIN analysis were achieved to test rhythmicity of expression. When only one was significant, the significance level is in lighter colour. Significance levels: ns: p>0.05; *: 0.05 ≥ p ≥ 0.01; **: 0.01 > p ≥ 0.001; ***: p <0.001

**Figure S9:**
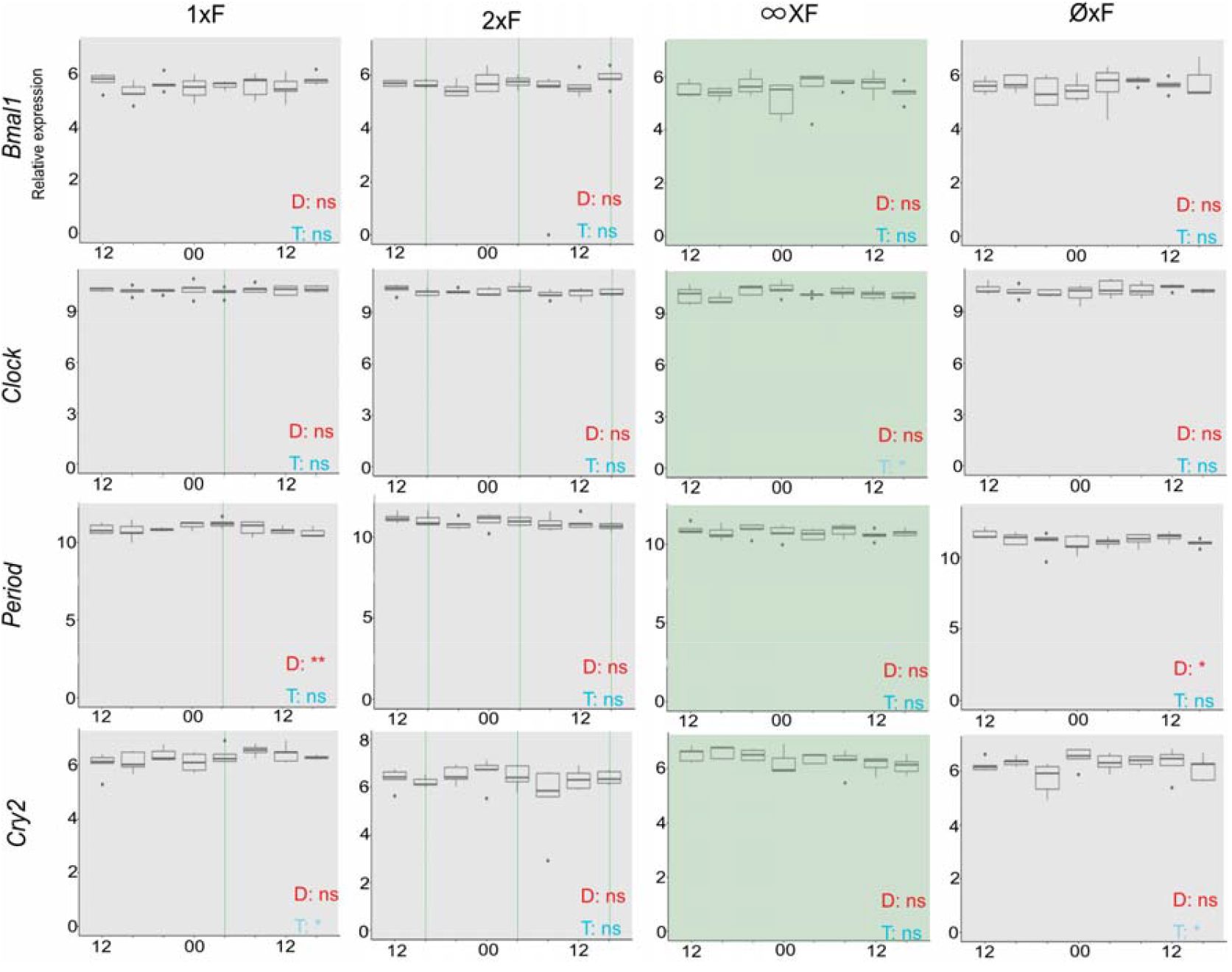
Core circadian clock gene expression under controlled condition of food availability. Food availability conditions tested were; fed once a day (1xF), fed twice a day (2xF), fed continuously (∞xF) and never fed (∅xF). Hours of feeding are indicated in green. Rhythmicity in expression were tested for the daily range of expression (D) and the tidal range (T). Cosinor adjustment and RAIN analysis were achieved to test rhythmicity of expression. When only one was significant, the significance level is in lighter colour. Significance levels: ns: p>0.05; *: 0.05 ≥ p ≥ 0.01; **: 0.01 > p ≥ 0.001; ***: p <0.001

**Figure S10:**
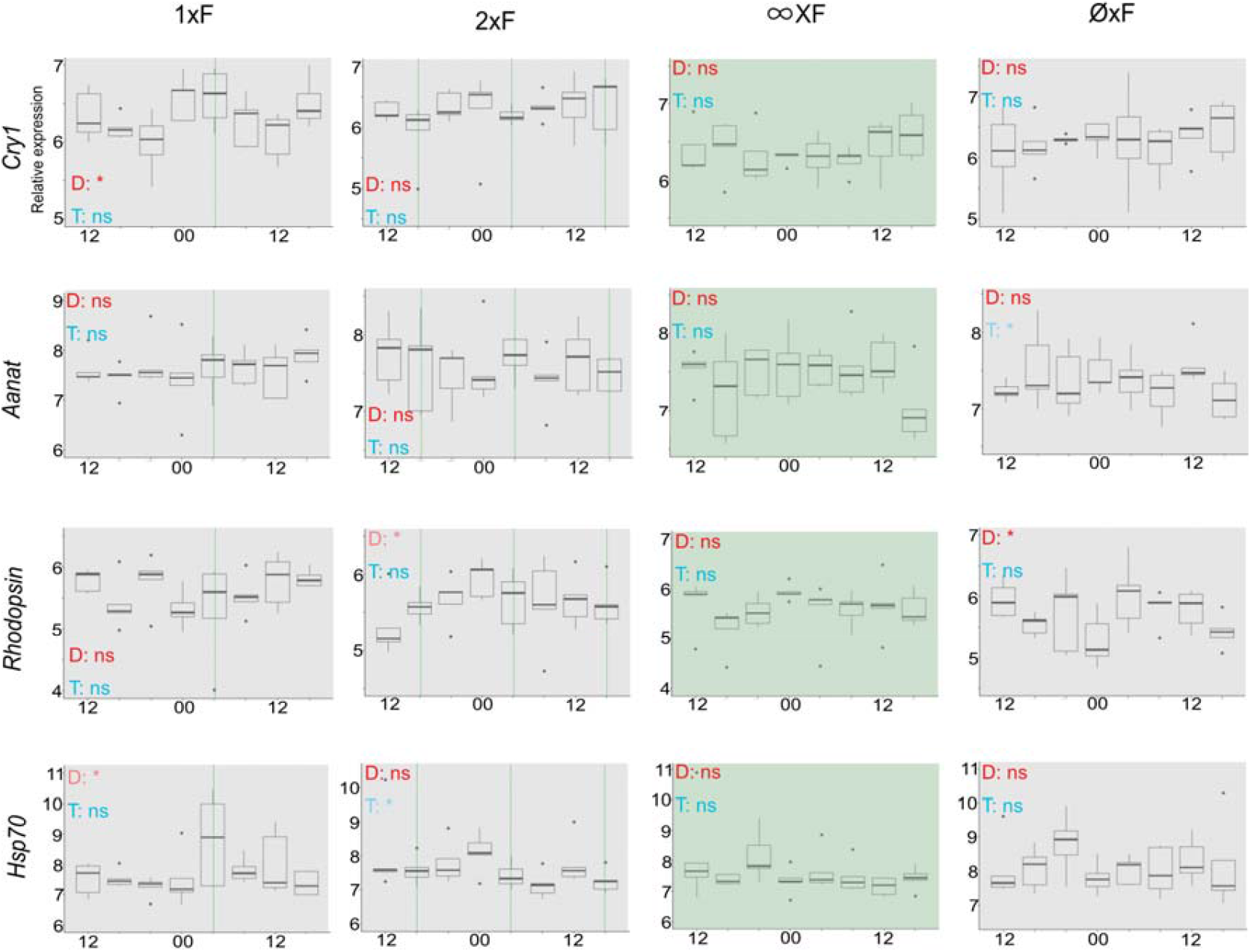
Expression of genes related to biological clock under controlled condition of food availability. Food availability conditions tested were; fed once a day (1xF), fed twice a day (2xF), fed continuously (∞xF) and never fed (∅xF). Hours of feeding are indicated in green. Rhythmicity in expression were tested for the daily range of expression (D) and the tidal range (T). Cosinor adjustment and RAIN analysis were achieved to test rhythmicity of expression. When only one was significant, the significance level is in lighter colour. Significance levels: ns: p>0.05; *: 0.05 ≥ p ≥ 0.01; **: 0.01 > p ≥ 0.001; ***: p <0.001

**Figure S11:**
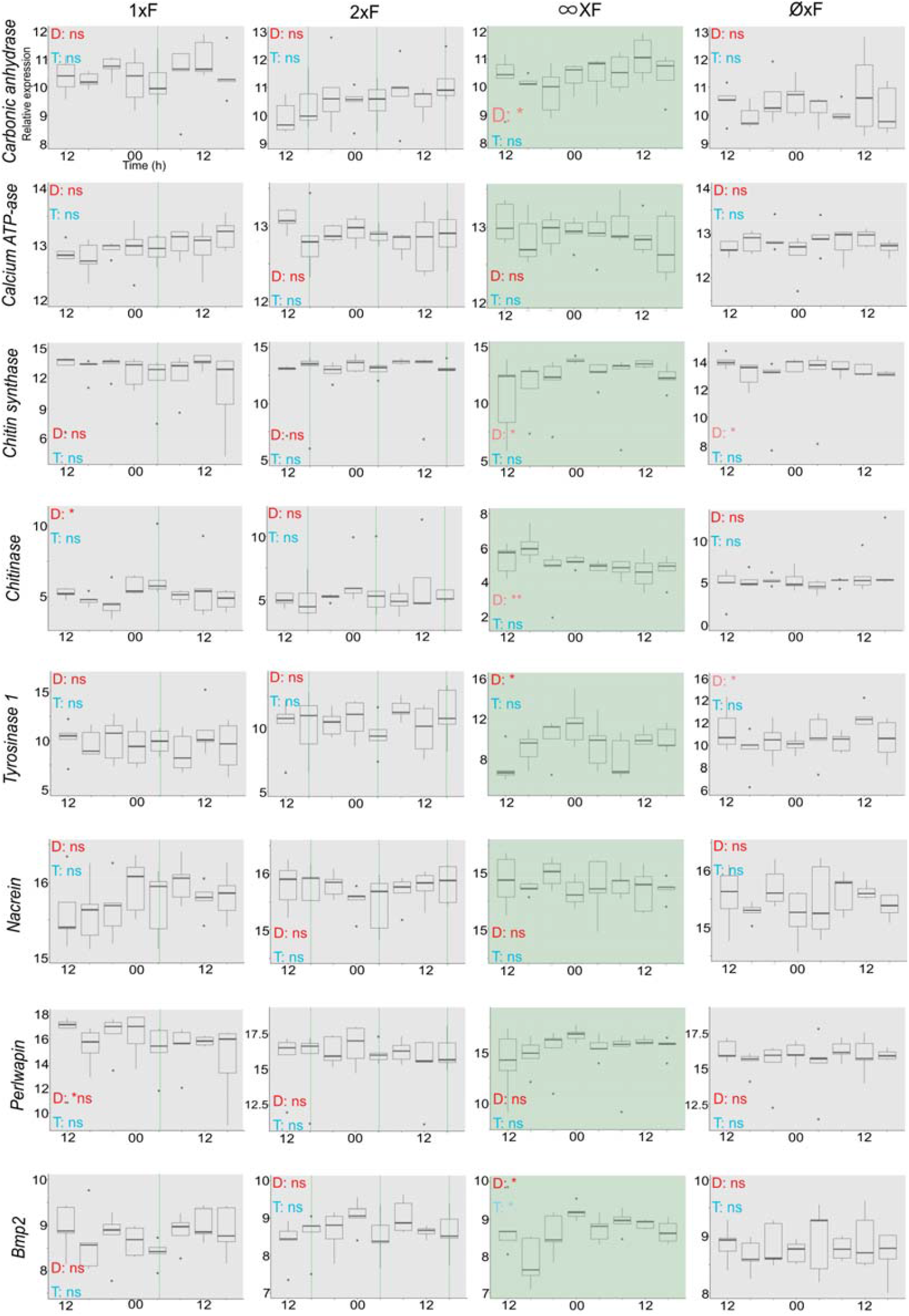
Biomineralisation related gene expression clock under controlled condition of food availability. Food availability conditions tested were; fed once a day (1xF), fed twice a day (2xF), fed continuously (∞xF) and never fed (∅xF). Hours of feeding are indicated in green. Rhythmicity in expression were tested for the daily range of expression (D) and the tidal range (T). Cosinor adjustment and RAIN analysis were achieved to test rhythmicity of expression. When only one was significant, the significance level is in lighter colour. Significance levels: ns: p>0.05; *: 0.05 ≥ p ≥ 0.01; **: 0.01 > p ≥ 0.001; ***: p <0.001

**Table S1:**
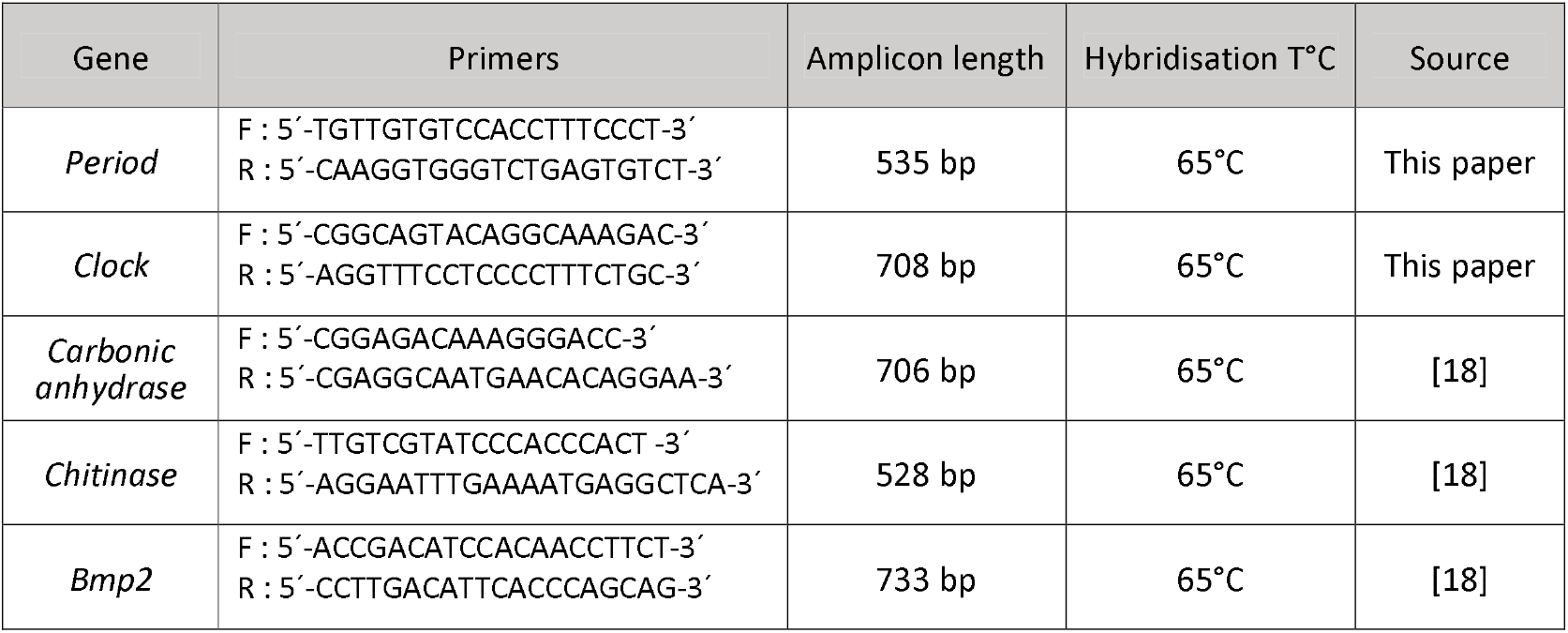
Targeted gene characteristics used in ISH on the posterior edge of the mantle of *M. galloprovincialis*.

**Table S2:** Statistics of circadian and circatidal periodicities at sea (SOLA). Bold values are significant. When Cosinor analysis as implemented in Discorhythm and RAIN analysis are both significant, the line is highlight in green. Grey numbers are outputs from RAIN analysis out of the framework of circatidal definition due to the sampling time interval.

|  | Circadian (24h+/-4h) |  |  |  | Circatidal (12h+/-2h) |  |  |  |
| --- | --- | --- | --- | --- | --- | --- | --- | --- |
|  | Cosinor |  | RAIN |  | Cosinor |  | RAIN |  |
|  | Period (h) | p-value | Period (h) | p-value | Period (h) | p-value | Period (h) | p-value |
| *bmal1* | 22 | 0.260 | 24 | 0.700 | 11 | 0.285 | 12 | 0.785 |
| *clock* | 20 | **0.001** | 20 | **5.2e-5** | 10 | 0.685 | 16 | 0.074 |
| *period* | 24 | **1e-24** | 24 | **1e-33** | 12 | **0.032** | 12 | **0.002** |
| *cry2* | 28 | 0.091 | 28 | 0.190 | 13 | 0.056 | 12 | 0.287 |
| *timeout* | NA | NA | NA | NA | NA | NA | NA | NA |
| *rorb* | NA | NA | NA | NA | NA | NA | NA | NA |
| *cry1* | 28 | 0.241 | 28 | **0.028** | 14 | **0.020** | 16 | **0.045** |
| *aanat* | 20 | 0.286 | 20 | 0.082 | 11 | 0.108 | 12 | 0.077 |
| *rhodopsin* | 26 | **0.042** | 24 | **0.018** | 13 | **0.050** | 12 | 0.103 |
| *hsp70* | 23 | **0.038** | 28 | **0.016** | 12 | 0.188 | 12 | 0.875 |

**Table S3:** Statistics of circadian and circatidal periodicities in photoperiodic (L:D 12:12) condition in aquaria. Bold values are significant. When Cosinor analysis as implemented in Discorhythm and RAIN analysis are both significant, the line is highlight in green. Grey numbers are outputs from RAIN analysis out of the framework of circatidal definition due to the sampling time interval.

|  | Circadian |  |  |  | Circatidal |  |  |  |
| --- | --- | --- | --- | --- | --- | --- | --- | --- |
|  | Cosinor |  | RAIN |  | Cosinor |  | RAIN |  |
|  | Period (h) | p-value | Period (h) | p-value | Period (h) | p-value | Period (h) | p-value |
| *bmal1* | 28 | 0.553 | 28 | 0.937 | 10 | 0.164 | 12 | 0.816 |
| *clock* | 27 | 0.082 | 24 | **0.042** | 12 | 0.426 | 12 | **0.869** |
| *period* | 20 | **0.047** | 24 | 0.107 | 14 | 0.110 | 16 | 0.156 |
| *cry2* | 28 | 0.193 | 28 | 0.249 | 14 | 0.120 | 16 | 0.424 |
| *timeout* | NA | NA | NA | NA | NA | NA | NA | NA |
| *rorb* | NA | NA | NA | NA | NA | NA | NA | NA |
| *cry1* | 20 | 0.406 | 24 | 0.729 | 10 | 0.162 | 8 | 0.725 |
| *aanat* | 28 | 0.335 | 28 | 0.581 | 10 | 0.274 | 8 | 0.175 |
| *rhodopsin* | 20 | 0.090 | 20 | 0.398 | 10 | 0.204 | 16 | **0.024** |
| *hsp70* | 28 | 0.906 | 28 | 0.752 | 12 | 0.391 | 8 | 0.824 |
| *carbonic anhydrase* | 23 | 0.187 | 24 | 0.391 | 11 | 0.917 | 12 | 0.792 |
| *Ca2+-ATPase* | 23 | 0.722 | 24 | 0.115 | 10 | 0.282 | 8 | 0.483 |
| *bmp2* | 20 | 0.750 | 28 | 0.908 | 14 | 0.482 | 16 | 0.527 |
| *chitin synthase* | 20 | 0.567 | 28 | 0.893 | 10 | 0.257 | 8 | 0.315 |
| *chitinase* | 20 | 0.226 | 28 | 0.948 | 14 | 0.511 | 12 | **0.447** |
| *nacrein* | 28 | 0.180 | 28 | 0.435 | 14 | 0.526 | 12 | 0.868 |
| *perlwapin* | 20 | 0.206 | 20 | 0.729 | 14 | 0.313 | 16 | **0.029** |
| *tyrosinase 1* | 20 | 0.361 | 20 | 0.819 | 10 | 0.257 | 16 | **0.044** |

**Table S4:** Statistics of circadian and circatidal periodicities in constant enlightenment (L:L) condition in aquaria. Bold values are significant. When Cosinor analysis as implemented in Discorhythm and RAIN analysis are both significant, the line is highlight in green. Grey numbers are outputs from RAIN analysis out of the framework of circatidal definition due to the sampling time interval.

|  | Circadian |  |  |  | Circatidal |  |  |  |
| --- | --- | --- | --- | --- | --- | --- | --- | --- |
|  | Cosinor |  | RAIN |  | Cosinor |  | RAIN |  |
|  | Period (h) | p-value | Period (h) | p-value | Period (h) | p-value | Period (h) | p-value |
| *bmal1* | 25 | 0.760 | 24 | 0.853 | 13 | 0.746 | 8 | 0.871 |
| *clock* | 22 | **3.9e-4** | 20 | **2.1e-4** | 10 | 0.132 | 16 | **0.007** |
| *period* | 21 | **0.043** | 24 | 0.073 | 11 | 0.403 | 16 | 0.720 |
| *cry2* | 24 | 0.059 | 28 | **0.014** | 11 | 0.087 | 12 | **0.298** |
| *timeout* | NA | NA | NA | NA | NA | NA | NA | NA |
| *rorb* | NA | NA | NA | NA | NA | NA | NA | NA |
| *cry1* | 20 | 0.327 | 20 | 0.881 | 14 | 0.356 | 16 | 0.640 |
| *aanat* | 20 | 0.583 | 28 | 0.994 | 12 | 0.076 | 16 | 0.126 |
| *rhodopsin* | 28 | 0.096 | 28 | **0.009** | 14 | 0.112 | 16 | 0.177 |
| *hsp70* | 28 | 0.177 | 24 | 0.186 | 11 | **0.022** | 12 | **0.157** |
| *carbonic anhydrase* | 27 | **0.025** | 24 | **0.005** | 13 | 0.352 | 12 | 0.737 |
| *Ca2+-ATPase* | 20 | 0.095 | 20 | 0.090 | 10 | **0.015** | 16 | 0.109 |
| *bmp2* | 20 | 0.056 | 20 | 0.185 | 10 | 0.132 | 16 | 0.311 |
| *chitin synthase* | 28 | 0.964 | 28 | 0.949 | 11 | 0.145 | 8 | 0.878 |
| *chitinase* | 20 | 0.076 | 20 | 0.256 | 14 | **0.015** | 16 | **0.004** |
| *nacrein* | 24 | **0.006** | 24 | **1.9e-5** | 12 | 0.240 | 16 | 0.088 |
| *perlwapin* | 28 | 0.860 | 24 | 0.896 | 11 | 0.195 | 8 | 0.814 |
| *tyrosinase 1* | 20 | 0.922 | 24 | 0.812 | 12 | 0.344 | 12 | 0.851 |

**Table S5:** Statistics of circadian and circatidal periodicities in constant darkness (D:D) condition in aquaria. Bold values are significant. When Cosinor analysis as implemented in Discorhythm and RAIN analysis are both significant, the line is highlight in green. Grey numbers are outputs from RAIN analysis out of the framework of circatidal definition due to the sampling time interval.

|  | Circadian |  |  |  | Circatidal |  |  |  |
| --- | --- | --- | --- | --- | --- | --- | --- | --- |
|  | Cosinor |  | RAIN |  | Cosinor |  | RAIN |  |
|  | Period (h) | p-value | Period (h) | p-value | Period (h) | p-value | Period (h) | p-value |
| *bmal1* | 23 | **0.011** | 20 | **0.016** | 10 | 0.138 | 16 | 0.377 |
| *clock* | 20 | **0.050** | 20 | 0.189 | 14 | **0.012** | 16 | **0.008** |
| *period* | 20 | 0.199 | 20 | 0.774 | 14 | 0.279 | 16 | 0.329 |
| *cry2* | 20 | 0.402 | 20 | 0.862 | 10 | 0.106 | 16 | 0.351 |
| *timeout* | NA | NA | NA | NA | NA | NA | NA | NA |
| *rorb* | NA | NA | NA | NA | NA | NA | NA | NA |
| *cry1* | 20 | 0.120 | 20 | 0.700 | 14 | 0.091 | 16 | **0.040** |
| *aanat* | 20 | 0.285 | 28 | 0.632 | 14 | 0.092 | 16 | **0.017** |
| *rhodopsin* | 20 | 0.089 | 20 | 0.156 | 14 | 0.376 | 16 | 0.165 |
| *hsp70* | 20 | 0.102 | 28 | **0.036** | 14 | **0.042** | 16 | **0.008** |
| *carbonic anhydrase* | 28 | 0.108 | 28 | **0.050** | 13 | 0.072 | 12 | 0.307 |
| *Ca2+-ATPase* | 20 | 0.165 | 28 | 0.258 | 14 | 0.252 | 16 | 0.170 |
| *bmp2* | 26 | 0.132 | 24 | 0.086 | 14 | 0.065 | 16 | **0.044** |
| *chitin synthase* | 21 | 0.214 | 28 | 0.460 | 14 | 0.107 | 12 | 0.242 |
| *chitinase* | 20 | 0.125 | 28 | 0.460 | 10 | **0.036** | 12 | 0.242 |
| *nacrein* | 28 | **0.031** | 28 | **0.030** | 10 | 0.292 | 16 | 0.593 |
| *perlwapin* | 26 | 0.272 | 28 | 0.351 | 13 | 0.146 | 12 | 0.435 |
| *tyrosinase 1* | 20 | 0.103 | 20 | 0.113 | 14 | **0.001** | 16 | **0.001** |

**Table S6:** Statistics of circadian and circatidal periodicities in no food availability (∅xF) and continuous darkness (D:D) condition in aquaria. Bold values are significant. When Cosinor analysis as implemented in Discorhythm and RAIN analysis are both significant, the line is highlight in green. Grey numbers are outputs from RAIN analysis out of the framework of circatidal definition due to the sampling time interval.

|  | Circadian |  |  |  | Circatidal |  |  |  |
| --- | --- | --- | --- | --- | --- | --- | --- | --- |
|  | Cosinor |  | RAIN |  | Cosinor |  | RAIN |  |
|  | Period (h) | p-value | Period (h) | p-value | Period (h) | p-value | Period (h) | p-value |
| *bmal1* | 28 | 0.412 | 24 | 0.921 | 14 | 0.630 | 16 | 0.718 |
| *clock* | 25 | 0.140 | 28 | 0.323 | 12 | 0.764 | 12 | 0.840 |
| *period* | 21 | **0.023** | 24 | **0.041** | 11 | 0.338 | 12 | 0.200 |
| *cry2* | 23 | 0.193 | 24 | 0.535 | 10 | **0.036** | 12 | 0.857 |
| *timeout* | NA | NA | NA | NA | NA | NA | NA | NA |
| *rorb* | NA | NA | NA | NA | NA | NA | NA | NA |
| *cry1* | 20 | 0.371 | 20 | 0.863 | 14 | 0.661 | 16 | 0.497 |
| *aanat* | 28 | 0.706 | 28 | 0.941 | 10 | **0.027** | 12 | 0.670 |
| *rhodopsin* | 21 | **0.034** | 20 | **0.020** | 14 | 0.534 | 16 | **0.006** |
| *hsp70* | 20 | 0.387 | 28 | 0.829 | 14 | 0.372 | 16 | 0.190 |
| *carbonic anhydrase* | 20 | 0.835 | 24 | 0.896 | 13 | 0.095 | 12 | 0.372 |
| *Ca2+-ATPase* | 20 | 0.564 | 28 | 0.900 | 10 | 0.143 | 16 | 0.183 |
| *bmp2* | 24 | 0.621 | 28 | 0.868 | 12 | 0.851 | 8 | 0.759 |
| *chitin synthase* | 23 | 0.451 | 20 | **0.016** | 10 | 0.126 | 16 | **0.039** |
| *chitinase* | 20 | 0.161 | 28 | 0.426 | 14 | 0.661 | 16 | 0.718 |
| *nacrein* | 20 | 0.578 | 24 | 0.642 | 12 | 0.188 | 12 | 0.120 |
| *perlwapin* | 27 | 0.536 | 28 | 0.905 | 10 | 0.195 | 12 | 0.794 |
| *tyrosinase 1* | 28 | 0.103 | 28 | **0.031** | 13 | 0.109 | 8 | 0.192 |

**Table S7:**
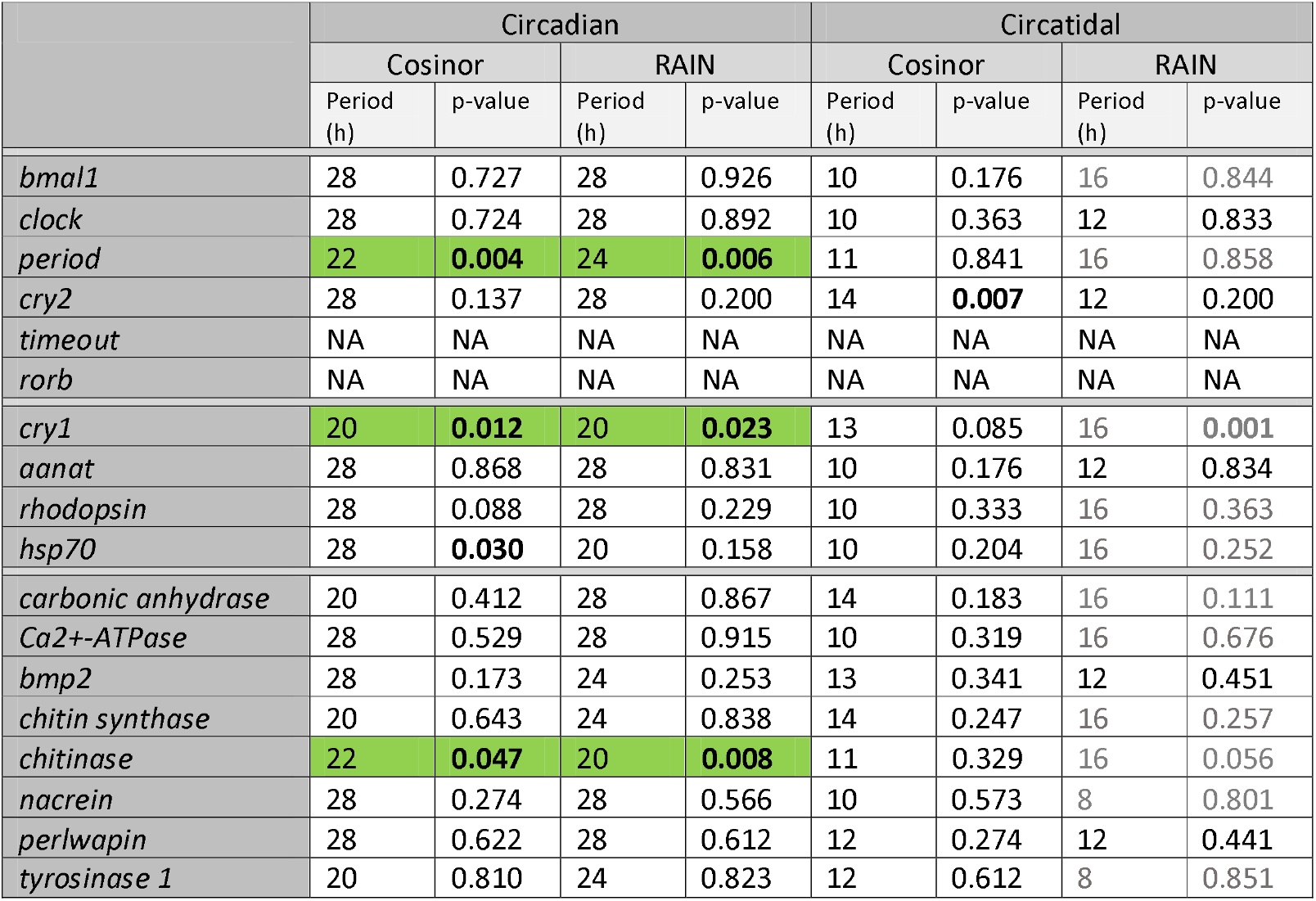
Statistics of circadian and circatidal periodicities in one feeding time per day (1xF) and continuous darkness (D:D) condition in aquaria. Bold values are significant. When Cosinor analysis as implemented in Discorhythm and RAIN analysis are both significant, the line is highlight in green. Grey numbers are outputs from RAIN analysis out of the framework of circatidal definition due to the sampling time interval.

**Table S8:**
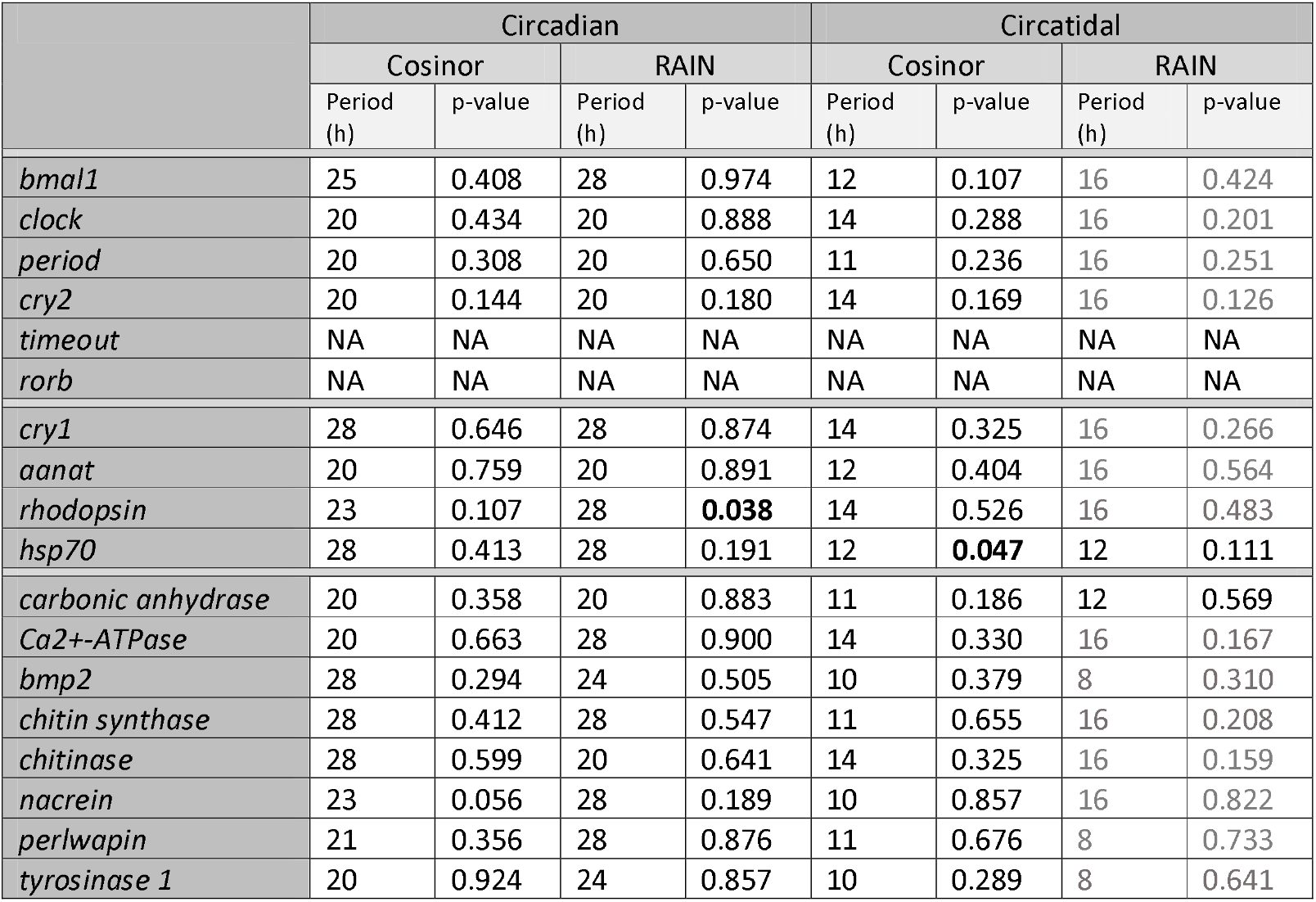
Statistics of circadian and circatidal periodicities in o two feeding time per day (2xF) and continuous darkness (D:D) condition in aquaria. Bold values are significant. When Cosinor analysis as implemented in Discorhythm and RAIN analysis are both significant, the line is highlight in green. Grey numbers are outputs from RAIN analysis out of the framework of circatidal definition due to the sampling time interval.

**Table S9:** Statistics of circadian and circatidal periodicities in continuous food availability (∞xF) and continuous darkness (D:D) condition in aquaria. Bold values are significant. When Cosinor analysis as implemented in Discorhythm and RAIN analysis are both significant, the line is highlight in green. Grey numbers are outputs from RAIN analysis out of the framework of circatidal definition due to the sampling time interval.

|  | Circadian |  |  |  | Circatidal |  |  |  |
| --- | --- | --- | --- | --- | --- | --- | --- | --- |
|  | Cosinor |  | RAIN |  | Cosinor |  | RAIN |  |
|  | Period (h) | p-value | Period (h) | p-value | Period (h) | p-value | Period (h) | p-value |
| *bmal1* | 20 | 0.405 | 28 | 0.203 | 13 | 0.263 | 16 | 0.519 |
| *clock* | 27 | 0.101 | 24 | 0.104 | 12 | **0.033** | 12 | 0.063 |
| *period* | 28 | 0.787 | 28 | 0.881 | 10 | 0.154 | 12 | 0.773 |
| *cry2* | 28 | 0.157 | 28 | 0.142 | 14 | 0.194 | 16 | 0.072 |
| *timeout* | NA | NA | NA | NA | NA | NA | NA | NA |
| *rorb* | NA | NA | NA | NA | NA | NA | NA | NA |
| *cry1* | 28 | 0.173 | 24 | 0.494 | 12 | 0.039 | 12 | 0.604 |
| *aanat* | 24 | 0.155 | 24 | 0.637 | 12 | 0.104 | 8 | 0.489 |
| *rhodopsin* | 20 | 0.324 | 20 | 0.123 | 13 | 0.082 | 16 | 0.084 |
| *hsp70* | 28 | 0.540 | 28 | 0.165 | 10 | 0.165 | 12 | 0.857 |
| *carbonic anhydrase* | 28 | 0.056 | 28 | **0.011** | 11 | 0.327 | 12 | 0.848 |
| *Ca2+-ATPase* | 20 | 0.316 | 24 | 0.791 | 11 | 0.201 | 12 | 0.882 |
| *bmp2* | 28 | **0.019** | 28 | **0.001** | 13 | **0.014** | 12 | 0.103 |
| *chitin synthase* | 28 | 0.313 | 28 | **0.019** | 14 | 0.334 | 12 | 0.092 |
| *chitinase* | 28 | 0.319 | 28 | **0.005** | 14 | 0.170 | 16 | 0.671 |
| *nacrein* | 28 | 0.858 | 28 | 0.823 | 11 | 0.297 | 12 | 0.733 |
| *perlwapin* | 20 | 0.202 | 20 | 0.328 | 14 | 0.298 | 16 | 0.129 |
| *tyrosinase 1* | 20 | **0.003** | 20 | **0.001** | 14 | 0.143 | 16 | **0.038** |

## References

1. Skinner HCW, Jahren AH. 2003 Biomineralization. In Trearise on Geochemistry, pp. 117–184. Elsevier Ltd. (doi:10.1016/B0-08-043751-6/08128-7)

2. Giuffre AJ, Hamm LM, Han N, De Yoreo JJ, Dove PM. 2013 Polysaccharide chemistry regulates kinetics of calcite nucleation through competition of interfacial energies. Proc. Natl. Acad. Sci. U. S. A. 110, 9261–9266. (doi:10.1073/pnas.1222162110)

3. Checa AG. 2018 Physical and biological determinants of the fabrication of Molluscan shell microstructures. Front. Mar. Sci. 5, 353. (doi:10.3389/fmars.2018.00353)

4. Schöne BR. 2008 The curse of physiology — challenges and opportunities in the interpretation of geochemical data from mollusk shells. Geo-Marine Letters 28, 269–285. (doi:10.1007/s00367-008-0114-6)

5. Louis V, Besseau L, Lartaud F. 2022 Step in Time: Biomineralisation of Bivalve’s Shell. Front. Mar. Sci. 9, 906085. (doi:10.3389/fmars.2022.906085)

6. Louis V, Desbordes F, Besseau L, Lartaud F. 2024 Bivalve shell growth from molecular to sclerochronological scale: Environment and intrinsic factors control increment deposition. Mar. Environ. Res. 202, 106730. (doi:10.1016/j.marenvres.2024.106730)

7. Richardson CA. 1988 Exogenous and endogenous rhythms of band formation in the shell of the clam Tapes philippinarum (Adams et Reeve, 1850). J. Exp. Mar. Biol. Ecol. 122, 105–126. (doi:10.1016/0022-0981(88)90179-7)

8. Richardson CA, Crisp DJ, Runham NW. 1980 An endogenous rhythm in shell deposition in Cerastoderma edule. Journal of the Marine Biological Association of the United Kingdom 60, 991–1004. (doi:10.1017/S0025315400042041)

9. Dunlap JC. 1999 Molecular bases for circadian clocks. Cell 96, 271–290. (doi:10.1016/s0092-8674(00)80566-8)

10. Rosbash M. 2009 The Implications of Multiple Circadian Clock Origins. PLoS Biol. 7, e1000062. (doi:10.1371/journal.pbio.1000062)

11. Aschoff J. 1981 Freerunning and Entrained Circadian Rhythms. In Biological Rhythms, pp. 81– 93. Boston, MA.: Springer US. (doi:10.1007/978-1-4615-6552-9_6)

12. Perrigault M, Tran D. 2017 Identification of the Molecular Clockwork of the Oyster Crassostrea gigas. PLoS One 12, e0169790. (doi:10.1371/journal.pone.0169790)

13. Jetten AM, Kurebayashi S, Ueda E. 2001 The ROR nuclear orphan receptor subfamily: Critical regulators of multiple biological processes. Prog. Nucleic Acid Res. Mol. Biol. 69, 205–247. (doi:10.1016/S0079-6603(01)69048-2)

14. Kwiatkowski ER, Schnytzer Y, Rosenthal JJC, Emery P. 2023 Behavioral circatidal rhythms require Bmal1 in Parhyale hawaiensis. Current Biology 33, 1867–1882. (doi:10.1016/j.cub.2023.03.015)

15. Zhang L, Green EW, Webster SG, Hastings Id MH, Wilcockson DC, Kyriacouid CP. 2023 The circadian clock gene bmal1 is necessary for co-ordinated circatidal rhythms in the marine isopod Eurydice pulchra (Leach). PLoS Genet. 19, e1011011. (doi:10.1371/journal.pgen.1011011)

16. Tran D, Perrigault M, Ciret P, Payton L. 2020 Bivalve mollusc circadian clock genes can run at tidal frequency. Proc. R. Soc. B 287, 20192440. (doi:10.1098/rspb.2019.2440)

17. Mat AM, Massabuau JC, Ciret P, Tran D. 2012 Evidence for a plastic dual circadian rhythm in the oyster Crassostrea gigas. Chronobiol. Int. 29, 857–867. (doi:10.3109/07420528.2012.699126)

18. Dunlap JC. 1999 Molecular bases for circadian clocks. Cell 96, 271–290. (doi:10.1016/s0092-8674(00)80566-8)

19. Stephan FK. 2002 The “Other” Circadian System: Food as a Zeitgeber. J. Biol. Rhythms 17, 284–292. (doi:10.1177/074873040201700402)

20. Bayne BL. 2004 Phenotypic Flexibility and Physiological Tradeoffs in the Feeding and Growth of Marine Bivalve Molluscs. Integr. Comp. Biol. 44, 425–432. (doi:10.1093/icb/44.6.425)

21. Tamayo D, Ibarrola I, Cigarría J, Navarro E. 2015 The effect of food conditioning on feeding and growth responses to variable rations in fast and slow growing spat of the Manila clam (Ruditapes philippinarum). J. Exp. Mar. Biol. Ecol. 471, 92–103. (doi:10.1016/j.jembe.2015.05.017)

22. SHOM. 2020 Horaires de marées gratuits du SHOM. See https://maree.shom.fr/harbor/BANYULS/hlt/0?date=2020-06-24&utc=standard (accessed on 13 September 2020).

23. Maire O, Amouroux JM, Duchêne JC, Grémare A. 2007 Relationship between filtration activity and food availability in the Mediterranean mussel Mytilus galloprovincialis. Mar. Biol. 152, 1293–1307. (doi:10.1007/s00227-007-0778-x)

24. Williams BG, Pilditch CA. 1997 The Entrainment of Persistent Tidal Rhythmicity in a Filter-Feeding Bivalve Using Cycles of Food Availability. J. Biol. Rhythms 12, 173–181. (doi:10.1177/074873049701200208)

25. Hall T. 2001 Biolign alignment and multiple contig editor.

26. Nei Masatoshi, Kumar S. 2000 Molecular evolution and phylogenetics. Oxford University Press.

27. Jones DT, Taylor WR, Thornton JM. 1992 The rapid generation of mutation data matrices from protein sequences. Bioinformatics 8, 275–282. (doi:10.1093/BIOINFORMATICS/8.3.275)

28. Kumar S, Stecher G, Li M, Knyaz C, Tamura K. 2018 MEGA X: Molecular Evolutionary Genetics Analysis across Computing Platforms. Mol. Biol. Evol. 35, 1547–1549. (doi:10.1093/molbev/msy096)

29. Gerdol M et al. 2020 Massive gene presence-absence variation shapes an open pan-genome in the Mediterranean mussel. Genome Biol. 21, 275. (doi:10.1186/s13059-020-02180-3)

30. Yang M, Derbyshire MK, Yamashita RA, Marchler-Bauer A. 2020 NCBI’s Conserved Domain Database and Tools for Protein Domain Analysis. Curr. Protoc. Bioinformatics 69, e90. (doi:10.1002/CPBI.90)

31. Mat AM, Perrigault M, Massabuau JC, Tran D. 2016 Role and expression of cry1 in the adductor muscle of the oyster Crassostrea gigas during daily and tidal valve activity rhythms. Chronobiol. Int. 33, 949–963. (doi:10.1080/07420528.2016.1181645)

32. Chaves I et al. 2011 The cryptochromes: Blue light photoreceptors in plants and animals. Annu. Rev. Plant Biol. 62, 335–364. (doi:10.1146/annurev-arplant-042110-103759)

33. Senthilan PR, Grebler R, Reinhard N, Rieger D, Helfrich-Förster C. 2019 Role of Rhodopsins as Circadian Photoreceptors in the Drosophila melanogaster. Biology. 8, 6. (doi:10.3390/BIOLOGY8010006)

34. Perrigault M, Andrade H, Bellec L, Ballantine C, Camus L, Tran D. 2020 Rhythms during the polar night: evidence of clock-gene oscillations in the Arctic scallop Chlamys islandica. Proc. Biol. Sci. 287, 20201001. (doi:10.1098/rspb.2020.1001)

35. Klein DC. 2007 Arylalkylamine N-Acetyltransferase: “the Timezyme”. Journal of Biological Chemistry 282, 4233–4237. (doi:10.1074/JBC.R600036200)

36. Zhu Y, Yu Z, Liao K, Zhang L, Ran Z, Xu J. 2022 Melatonin in razor clam Sinonovacula constrictal: Examination of metabolic pathways, tissue distribution, and daily rhythms. Aquaculture 560, 738548. (doi:10.1016/j.aquaculture.2022.738548)

37. Sleight VA, Antczak P, Falciani F, Clark MS, Cowen L. 2020 Computationally predicted gene regulatory networks in molluscan biomineralization identify extracellular matrix production and ion transportation pathways. Bioinformatics 36, 1326–1332. (doi:10.1093/bioinformatics/btz754)

38. Toyohara H, Hosoi M, Hayashi I, Kubota S, Hashimoto H, Yokoyama Y. 2005 Expression of HSP70 in response to heat-shock and its cDNA cloning from Mediterranean blue mussel. Fisheries Science 2005 71:2 71, 327–332. (doi:10.1111/J.1444-2906.2005.00968.X)

39. Anestis A, Pörtner HO, Michaelidis B. 2010 Anaerobic metabolic patterns related to stress responses in hypoxia exposed mussels Mytilus galloprovincialis. J. Exp. Mar. Biol. Ecol. 394, 123–133. (doi:10.1016/J.JEMBE.2010.08.008)

40. Engel J. 2017 Chapter 6: Formation of Mollusk Shells. In A Critical Survey of Biomineralization (ed J Engel), pp. 29–40. (doi:10.1007/978-3-319-47711-4)

41. Miglioli A, Dumollard R, Balbi T, Besnardeau L, Canesi L. 2019 Characterization of the main steps in first shell formation in Mytilus galloprovincialis: Possible role of tyrosinase. Proceedings of the Royal Society B: Biological Sciences 286, 20192043. (doi:10.1098/rspb.2019.2043)

42. Treccani L, Mann K, Heinemann F, Fritz M. 2006 Perlwapin, an Abalone Nacre Protein with Three Four-Disulfide Core (Whey Acidic Protein) Domains, Inhibits the Growth of Calcium Carbonate Crystals. Biophys. J. 91, 2601–2608. (doi:10.1529/BIOPHYSJ.106.086108)

43. Song X, Liu Z, Wang L, Song L. 2019 Recent advances of shell matrix proteins and cellular orchestration in marine molluscan shell biomineralization. Front. Mar. Sci. 6, 41. (doi:10.3389/fmars.2019.00041)

44. Miyashita T, Hanashita T, Toriyama M, Takagi R, Akashika T, Higashikubo N. 2008 Gene cloning and biochemical characterization of the BMP-2 of Pinctada fucata. Biosci. Biotechnol. Biochem. 72, 37–47. (doi:10.1271/bbb.70302)

45. Zhao M, Shi Y, He M, Huang X, Wang Q. 2016 PfSMAD4 plays a role in biomineralization and can transduce bone morphogenetic protein-2 signals in the pearl oyster Pinctada fucata. BMC Dev. Biol. 16, 9. (doi:10.1186/s12861-016-0110-4)

46. Besseau L, Benyassi A, Møller M, Coon SL, Weller JL, Boeuf G, Klein DC, Falcón J. 2006 Melatonin pathway: breaking the ‘high-at-night’ rule in trout retina. Exp. Eye Res. 82, 620– 627. (doi:10.1016/J.EXER.2005.08.025)

47. NanoString Technologies Inc. 2017 Gene Expression Data Analysis Guidelines.

48. Comeau LA, Babarro JMF, Longa A, Padin XA. 2018 Valve-gaping behavior of raft-cultivated mussels in the Ría de Arousa, Spain. Aquac. Rep. 9, 68–73. (doi:10.1016/J.AQREP.2017.12.005)

49. Tran D, Ciret P, Ciutat A, Durrieu G, Massabuau J-C. 2003 Estimation of potential and limits of bivalve closure response to detect contaminants: Application to cadmium. Environ. Toxicol. Chem. 22, 914–920. (doi:10.1002/etc.5620220432)

50. Comeau LA, Babarro JMF, Longa A, Padin XA. 2018 Valve-gaping behavior of raft-cultivated mussels in the Ría de Arousa, Spain. Aquac. Rep. 9, 68–73. (doi:10.1016/J.AQREP.2017.12.005)

51. Bertolini C, Rubinetti S, Umgiesser G, Witbaard R, Bouma TJ, Rubino A, Pastres R. 2021 How to cope in heterogeneous coastal environments: Spatio-temporally endogenous circadian rhythm of valve gaping by mussels. Science of the Total Environment 768, 145085. (doi:10.1016/j.scitotenv.2021.145085)

52. Carlucci M, Krisciunas A, Li H, Gibas P, Koncevicius K, Petronis A, Oh G. 2020 DiscoRhythm: an easy-to-use web application and R package for discovering rhythmicity. Bioinformatics 36, 1952–1954. (doi:10.1093/BIOINFORMATICS)

53. Thaben PF, Westermark PO. 2014 Detecting rhythms in time series with RAIN. J. Biol. Rhythms 29, 391–400. (doi:10.1177/0748730414553029)

54. R Core Team. 2020 R: A Language and Environment for Statistical Computing.

55. Benjamini Y, Hochberg Y. 2000 On the Adaptive Control of the False Discovery Rate in Multiple Testing With Independent Statistics: Journal of Educational and behavioral Statistics 25, 60– 83. (doi:10.3102/10769986025001060)

56. Schmid B, Helfrich-Förster C, Yoshii T. 2011 A new ImageJ plugin ‘ActogramJ’ for chronobiological analyses. J. Biol. Rhythms 26, 464–467.

57. Gouthiere L, Claustrat B, Brun J, Mauvieux B. 2005 Éléments méthodologiques complémentaires dans l’analyse des rythmes : Recherche de périodes, modélisation. Exemples de la Mélatonine plasmatique et de courbes de températures. Pathologie Biologie 53, 285–289. (doi:10.1016/j.patbio.2004.12.025)

58. Gouthiere L, Mauvieux B, Davenne D, Waterhouse J. 2005 Complementary methodology in the analysis of rhythmic data, using examples from a complex situation, the rhythmicity of temperature in night shift workers. Biol. Rhythm Res. 36, 177–193. (doi:10.1080/09291010400026298)

59. Sachs M. 2014 cosinor: Tools for estimating and predicting the cosinor model. 60.

60. Shah AS. 2020 card: Cardiovascular and Autonomic Research Design.

61. Cornelissen G. 2014 Cosinor-based rhythmometry. Theor. Biol. Med. Model. 11, 1–24. (doi:10.1186/1742-4682-11-16)

62. Chapman EC, O’Dell AR, Meligi NM, Parsons DR, Rotchell JM. 2017 Seasonal expression patterns of clock-associated genes in the blue mussel Mytilus edulis. Chronobiol. Int. 34, 1300–1314. (doi:10.1080/07420528.2017.1363224)

63. Mat AM et al. 2020 Biological rhythms in the deep-sea hydrothermal mussel Bathymodiolus azoricus. Nat. Commun. 11, 3454. (doi:10.1038/s41467-020-17284-4)

64. Liu Y, He Q, Yao H, Lin Z, Dong Y. 2022 Circadian clock genes Bmal1 and Period may regulate nocturnal spawning by controlling sex hormone secretion in razor clam Sinonovacula constricta. Front. Mar. Sci. 9, 1–13. (doi:10.3389/fmars.2022.1074816)

65. Zhu H, Sauman I, Yuan Q, Casselman A, Emery-Le M, Emery P, Reppert SM. 2008 Cryptochromes define a novel circadian clock mechanism in monarch butterflies that may underlie sun compass navigation. PLoS Biol. 6, 0138–0155. (doi:10.1371/journal.pbio.0060004)

66. Rock A, Wilcockson D, Last KS. 2022 Towards an Understanding of Circatidal Clocks. Front. Physiol. 13, 1–7. (doi:10.3389/fphys.2022.830107)

67. Takekata H, Matsuura Y, Goto SG, Satoh A, Numata H. 2012 RNAi of the circadian clock gene period disrupts the circadian rhythm but not the circatidal rhythm in the mangrove cricket. Biol. Lett. 8, 488–491. (doi:10.1098/RSBL.2012.0079)

68. Takekata H, Numata H, Shiga S, Goto SG. 2014 Silencing the circadian clock gene Clock using RNAi reveals dissociation of the circatidal clock from the circadian clock in the mangrove cricket. J. Insect Physiol. 68, 16–22. (doi:10.1016/J.JINSPHYS.2014.06.012)

69. Zhang L, Hastings MH, Green EW, Tauber E, Sladek M, Webster SG, Kyriacou CP, Wilcockson DC. 2013 Dissociation of Circadian and Circatidal Timekeeping in the Marine Crustacean Eurydice pulchra. Current Biology 23, 1863–1873. (doi:10.1016/J.CUB.2013.08.038)

70. Bulla M, Oudman T, Bijleveld AI, Piersma T, Kyriacou CP. 2017 Marine biorhythms: Bridging chronobiology and ecology. Philosophical Transactions of the Royal Society B: Biological Sciences 372, 20160253. (doi:10.1098/rstb.2016.0253)

71. Bartness TJ, Albers HE. 2000 Activity Patterns and the Biological Clock in Mammals. In Activity patterns in small mammals, pp. 23–47. Springer, Berlin, Heidelberg. (doi:10.1007/978-3-642-18264-8_3)

72. Gosling E. 2004 Bivalve Molluscs: biology, ecology and culture. Oxford and Malden (Massachusetts). (doi:10.1002/9780470995532)

73. Richardson CA, Runham NW, Crisp DJ. 1981 A histological and ultrastructural study of the cells of the mantle edge of a marine bivalve, Cerastoderma edule. Tissue Cell 13, 715–730. (doi:10.1016/S0040-8166(81)80008-0)

74. Okada Y, Yamaura K, Suzuki T, Itoh N, Osada M, Takahashi KG. 2013 Molecular characterization and expression analysis of chitinase from the Pacific oyster Crassostrea gigas. Comparative Biochemistry and Physiology-Part B 165, 83–89. (doi:10.1016/j.cbpb.2013.03.008)

75. Ivanina A V., Falfushynska HI, Beniash E, Piontkivska H, Sokolova IM. 2017 Biomineralization-related specialization of hemocytes and mantle tissues of the Pacific oyster Crassostrea gigas. Journal of Experimental Biology 220, 3209–3221. (doi:10.1242/jeb.160861)

76. Mount AS, Wheeler AP, Paradkar RP, Snider D. 2004 Hemocyte-Mediated Shell Mineralization in the Eastern Oyster. Science. 304, 297–300.

77. Eggermont M et al. 2020 The blue mussel inside: 3D visualization and description of the vascular-related anatomy of Mytilus edulis to unravel hemolymph extraction. Sci. Rep. 10, 1– 16. (doi:10.1038/s41598-020-62933-9)

78. Tessmar-Raible K, Raible F, Arboleda E. 2011 Another place, another timer: Marine species and the rhythms of life. BioEssays 33, 165–172. (doi:10.1002/bies.201000096)

79. Mrosovsky N. 1999 Masking: history, definitions, and measurement. Chronobiol. Int. 16, 415– 429. (doi:10.3109/07420529908998717)

80. Oliphant A, Sia CY, Kyriacou CP, Wilcockson DC, Hastings MH. 2025 Expression of clock genes tracks daily and tidal time in brains of intertidal crustaceans Eurydice pulchra and Parhyale hawaiensis. Current Biology 12, 2802–2815. (doi:10.1016/j.cub.2025.04.047)

81. Wang X et al. 2013 Oyster Shell Proteins Originate from Multiple Organs and Their Probable Transport Pathway to the Shell Formation Front. PLoS One 8, e66522. (doi:10.1371/journal.pone.0066522)

82. Li S, Liu Y, Liu C, Huang J, Zheng G, Xie L, Zhang R. 2016 Hemocytes participate in calcium carbonate crystal formation, transportation and shell regeneration in the pearl oyster Pinctada fucata. Fish Shellfish Immunol. 51, 263–270. (doi:10.1016/J.FSI.2016.02.027)

83. Mat AM, Charles J, Ciret P, Tran D. 2014 Looking for the clock mechanism responsible for circatidal behavior in the oyster Crassostrea gigas. Mar. Biol. 161, 89–99. (doi:10.1007/s00227-013-2317-2)

84. Lutz RA, Rhoads DC. 1977 Anaerobiosis and a Theory of Growth Line Formation. Science 198, 1222–1227. (doi:10.1126/science.198.4323.1222)

85. Botté A, Payton L, Tran D. 2023 Artificial light at night at environmental intensities disrupts daily rhythm of the oyster Crassostrea gigas. Mar. Pollut. Bull. 191, 114850. (doi:10.1016/J.MARPOLBUL.2023.114850)

86. Duarte C et al. 2019 Artificial light pollution at night (ALAN) disrupts the distribution and circadian rhythm of a sandy beach isopod. Environmental Pollution 248, 565–573 (doi:10.1016/j.envpol.2019.02.037)

